# Psychiatric risk implications of adolescent exposure to environmental insecticides: a systematic review of rodent studies

**DOI:** 10.1101/2025.11.06.687006

**Authors:** Michelle X. Chen, Benjamin Hing, Robert J. Taylor, Hanna E. Stevens

## Abstract

Adolescence is a sensitive period of neurodevelopment marked by remodeling of brain circuits that support cognitive development and emotion and behavior regulation. These maturation processes heighten psychiatric vulnerability to environmental exposures, including to toxicants such as insecticides. Epidemiological studies show widespread adolescent insecticide exposure and increasingly link this with psychiatric outcomes, yet underlying neural mechanisms remain poorly understood. Preclinical studies can clarify these associations and identify insecticide-induced mechanisms that may disrupt neurodevelopment and produce consequent long-term behavioral outcomes. Here, we performed a systematic review of rodent studies following PRISMA guidelines. 50 original articles met inclusion criteria, examining neurotoxic outcomes following insecticide exposure during adolescence (postnatal days 21-60). Outcomes were categorized into four domains: neurocognitive, neuropsychiatric, neurobiological, and general neurotoxicity. Risk of bias was assessed using the SYRCLE Risk of Bias tool. Across studies, insecticide exposure during adolescence led to learning and memory impairments and tended to increase depression relevant behaviors, alter locomotor activity, and produce general neurotoxic effects. Mechanistic findings highlighted disruptions in cholinergic and monoaminergic signaling, oxidative stress, neuroimmune changes, and cell death and other neurodegenerative processes. Together, these findings indicate adolescent insecticide exposure disrupts multiple neural systems with behavioral consequences relevant to adolescent development and psychiatric risk. Future research should model real-world exposures (e.g. dose, timing) to better inform translational understanding of adolescent psychiatric vulnerability. Because many life-long neuropsychiatric disorders emerge in adolescence, identifying how modifiable environmental exposures shape risk offers an opportunity for prevention and intervention strategies to alter the course of disease across the lifespan.

## 1. Introduction

Neuropsychiatric disorders greatly impact those living with these conditions, oftentimes with challenges in cognition, emotion regulation, and behaviors. The prevalence of these brain disorders has risen in recent decades, particularly in children and adolescents (1–3). While this increase in part reflects improved screening and diagnostics, it also highlights an urgency to understand contributors to disease risk. Recently, there has been increased attention toward identifying environmental contributors to declining youth mental health, including environmental toxicants (4, 5). Adolescence is a neurodevelopmental period when cortical and mesolimbic circuits mature (6–8), making the adolescent brain particularly sensitive to external stressor effects, including toxicants. This neurodevelopmental period is particularly relevant to the risk for neuropsychiatric outcomes such as depression, which implicate many shared brain circuits, frequently emerge during adolescence, and oftentimes persist through adulthood (8–11).

Insecticides, compounds used to kill or repel insects, are one category of environmental toxicants. These compounds, which have mostly been designed to target insect nervous systems, are used in agriculture for crop management and in homes for insect repellent. Common human exposure routes include ingestion of food with insecticide residues, dermal or inhaled contact during household or agricultural use, and spray drift from nearby applications (12). Insecticide exposure is widespread, including for adolescents, with metabolites detected in 90-100% of individuals worldwide (13–15). This pervasive exposure underscores the critical need to understand their neurotoxic effects. There are currently 20 insecticide classes, five of which are most commonly used, with primary neurotoxic targets associated with brain outcomes in children and adolescents, including impairments with cognition, behavioral challenges, and psychiatric symptoms ((16), Table 1). Most epidemiological studies examine gestational or early childhood insecticide exposures linked to brain changes and increased risk for neurodevelopmental disorders (17–19). Emerging human evidence also suggests adolescence is a vulnerable period to insecticides, with exposures correlating with cognitive and behavioral symptoms in teenagers (20–28). A gap remains in understanding whether adolescent insecticide exposure directly causes these neuropsychiatric symptoms and, if so, mechanisms by which this occurs, limiting prevention and intervention strategies for youth at risk.

**Table 1.**
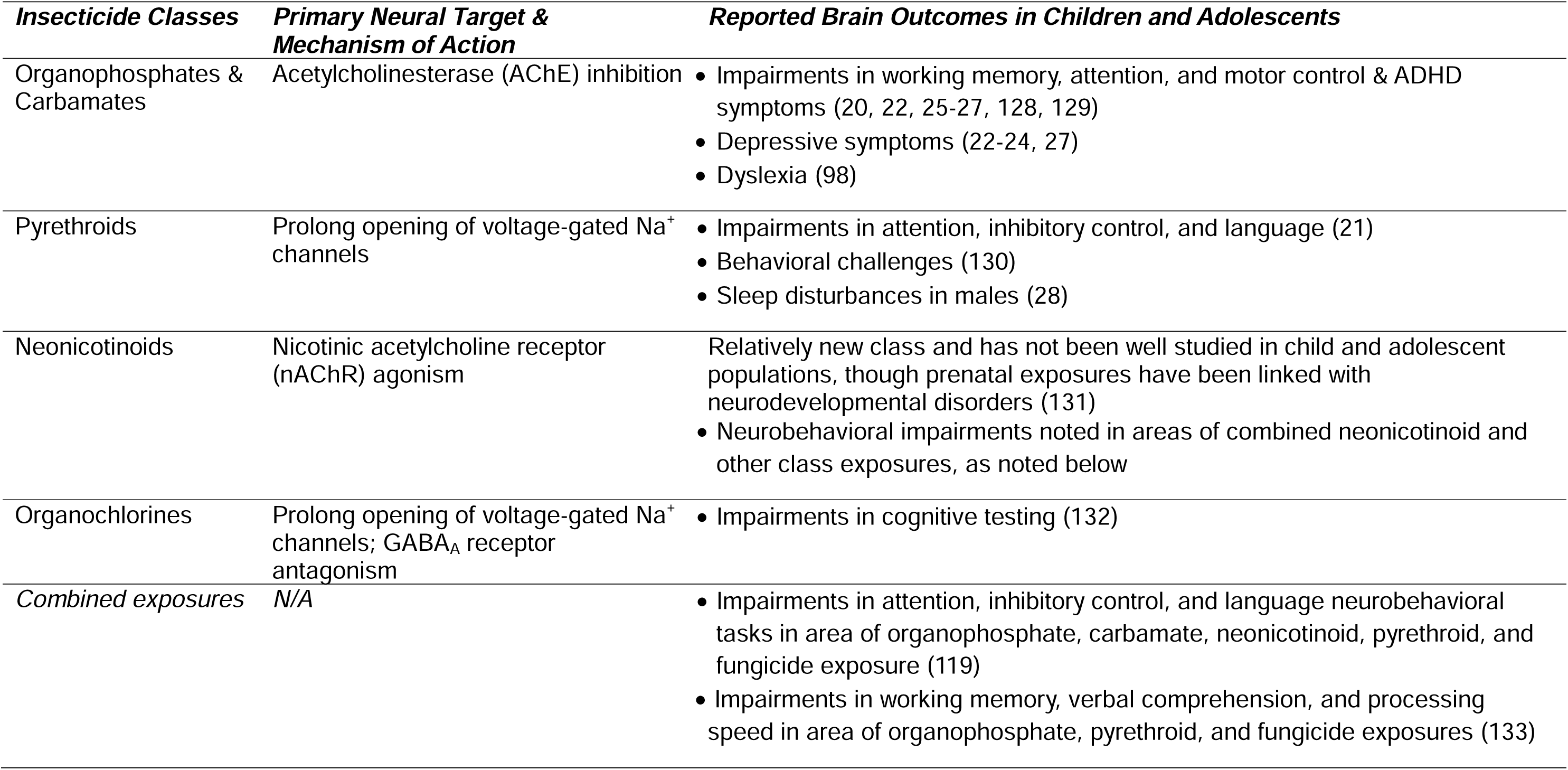
Neurotoxic mechanisms of frequently used insecticide classes and reported child and adolescent brain outcomes from occupational and environmental exposures.

| ***Insecticide Classes*** | ***Primary Neural Target & Mechanism of Action*** | ***Reported Brain Outcomes in Children and Adolescents*** |
| --- | --- | --- |
| Organophosphates & Carbamates | Acetylcholinesterase (AChE) inhibition | • Impairments in working memory, attention, and motor control & ADHD symptoms (20, 22, 25-27, 128, 129)<br>• Depressive symptoms (22-24, 27)<br>• Dyslexia (98) |
| Pyrethroids | Prolong opening of voltage-gated Na<sup>+</sup> channels | • Impairments in attention, inhibitory control, and language (21)<br>• Behavioral challenges (130)<br>• Sleep disturbances in males (28) |
| Neonicotinoids | Nicotinic acetylcholine receptor (nAChR) agonism | Relatively new class and has not been well studied in child and adolescent populations, though prenatal exposures have been linked with neurodevelopmental disorders (131)<br>• Neurobehavioral impairments noted in areas of combined neonicotinoid and other class exposures, as noted below |
| Organochlorines | Prolong opening of voltage-gated Na<sup>+</sup> channels; GABA<sub>A</sub> receptor antagonism | • Impairments in cognitive testing (132) |
| *Combined exposures* | *N/A* | • Impairments in attention, inhibitory control, and language neurobehavioral tasks in area of organophosphate, carbamate, neonicotinoid, pyrethroid, and fungicide exposure (119)<br>• Impairments in working memory, verbal comprehension, and processing speed in area of organophosphate, pyrethroid, and fungicide exposures (133) |

Animal models are essential for studying neurotoxic impacts and insecticide mechanisms. Rats and mice are especially valuable for modeling neurodevelopment because they share similar cellular programming with humans and undergo comparable brain maturation patterns (29, 30), (Figure 1). While *in vitro* models are valuable for identifying toxicity mechanisms in specific cell types, they are limited by absence of integrated whole-body physiology. In contrast, whole animals incorporate detoxification and integrative biological systems as occurs in humans, underscoring their relevance in neurotoxicology research. As in epidemiology, animal models have provided valuable insights regarding the effects of insecticides during gestational and neonatal development (31, 32). However, a synthesis of studies examining exposures during adolescence has not yet been done. Therefore, to address this gap, we conducted a systematic review of preclinical rodent studies that investigated neurotoxic effects of adolescent insecticide exposure. Broadening our understanding of how insecticide exposure impacts all neurodevelopmental periods is essential to inform public health policies and screening practices to protect young people.

**Figure 1.**
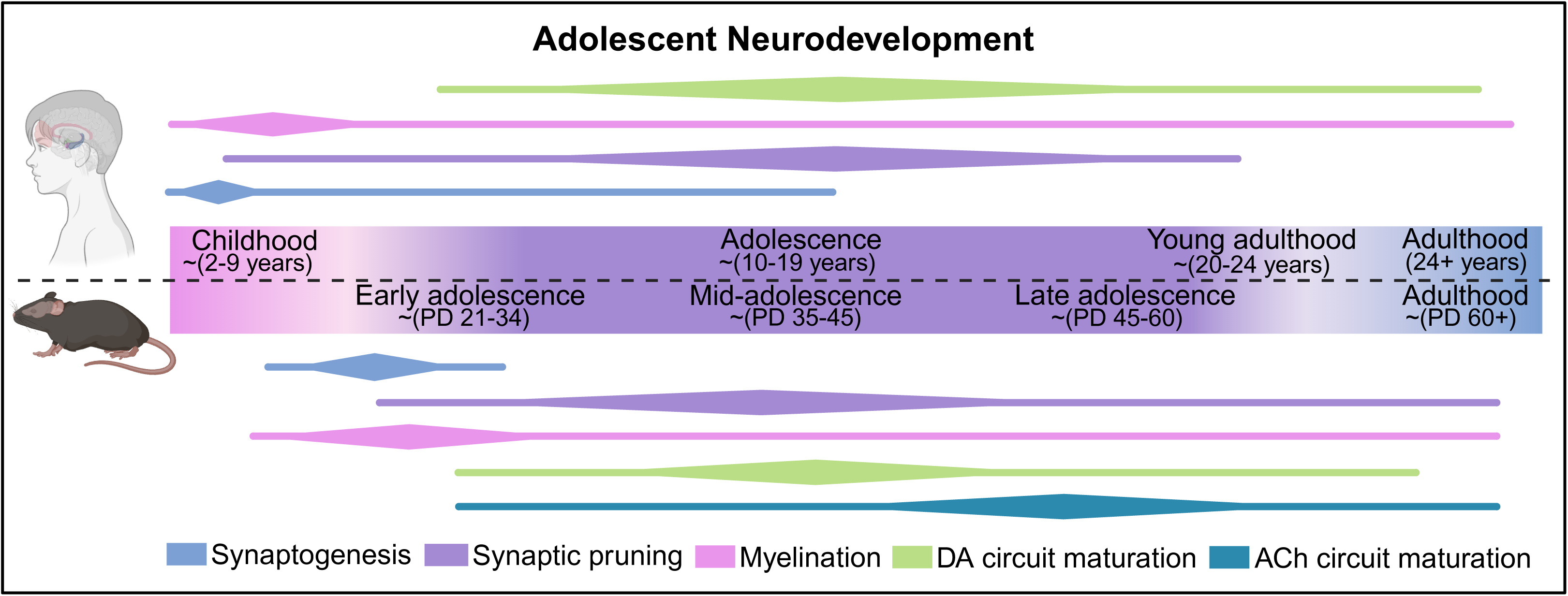
Adolescent neurodevelopmental processes in rodents and humans. Bars represent approximate timing of synaptogenesis, synaptic pruning, myelination, dopaminergic (DA) circuit maturation, cholinergic (ACh) circuit maturation, with peak activity indicated by the widest part of the diamond. No studies on human cholinergic circuit maturation performed to date, so ACh circuit maturation bar was not included for human development. Adapted from (8, 11, 29, 30, 127). *Made with BioRender*.

## 2. Methods

### Search Strategy & Screening

This systematic review followed PRISMA (Preferred Reporting Items for Systematic Reviews and Meta-Analyses) guidelines (33). The objective was to identify rat and mouse studies investigating neurotoxic effects of insecticide exposure during adolescence. While puberty can be defined as the period during which sexual maturity is reached, “adolescence” has been more difficult to define in rodents, as “puberty” and “adolescence” are not interchangeable terms. However, in the broadest sense, it has been suggested age of rodent weaning (postnatal day 21, PD21) until young adulthood (PD60) can be considered “adolescence,” as brain development during this period parallels that of human adolescent development, approximately ages 10 to 19 (Figure 1), (8, 34–36).

Literature searches were conducted in Embase, PubMed, and Scopus using Boolean terms (*Supplementary Materials).* The initial search (July 10, 2024) identified 4024 citations, and an update (August 15, 2025) identified 234 additional citations. Duplicates were removed following the Bramer method (37), leaving 2989 unique citations. Titles and abstracts were screened in Rayyan (Cambridge, MA) by two independent reviewers using predetermined exclusion criteria:

i. conference related; (ii) no neurotoxic outcome studied; (iii) no insecticide studied; (iv) study not performed in rats or mice.

### Eligibility & Extraction

Full-text screening was applied to 507 articles by two independent reviewers, of which 50 met inclusion criteria: (i) full-text journal articles; (ii) primary literature; (iii) conducted in rats or mice; (iv) exposure occurred only between PD21-60; (v): investigated an insecticide, (vi): investigated a neurotoxic outcome. Articles were excluded if they: (i) were review papers; (ii) did not study neurotoxic outcome; (iii) exposure occurred outside of the PD21-60 timepoint; (iv): investigated only a non-insecticide toxin (including other categories of pesticides, e.g. herbicides); (v): only *in vitro* studies; (vi) only ADME articles (vii) were published in languages other than English.

Data were extracted by one researcher focusing on comparisons between control and insecticide-only groups for nervous system-related outcomes. We recorded pre-specified information: publication characteristics, experimental model, exposure methods, and neurotoxic outcomes. Outcomes were categorized into four domains (neurocognitive, neuropsychiatric, neurobiological, general neurotoxic) and organized by insecticide class (Table 1). Articles reporting multiple relevant outcomes contributed more than one study (e.g., short- and long-term assessments were delineated separately). Full study-level details are provided (*Supplementary Tables)*.

### Quality Assessment

Risk of bias was evaluated using SYRCLE’s risk of bias tool (38), rating each study as high, low, or unclear risk across ten domains: 1) Random sequence generation, 2) Comparable baseline characteristics, 3) Allocation concealment, 4) Randomized housing, 5) Experimenter blinded to drug treatment, 6) Randomized animal for outcome assessment, 7) Outcome assessor blinded to drug treatment, 8) Incomplete outcome data reported, 9) No selective outcome reported, 10) Other sources of bias addressed.

## 3. Results

50 studies matched inclusion criteria and were included (Figure 2).

**Figure 2.**
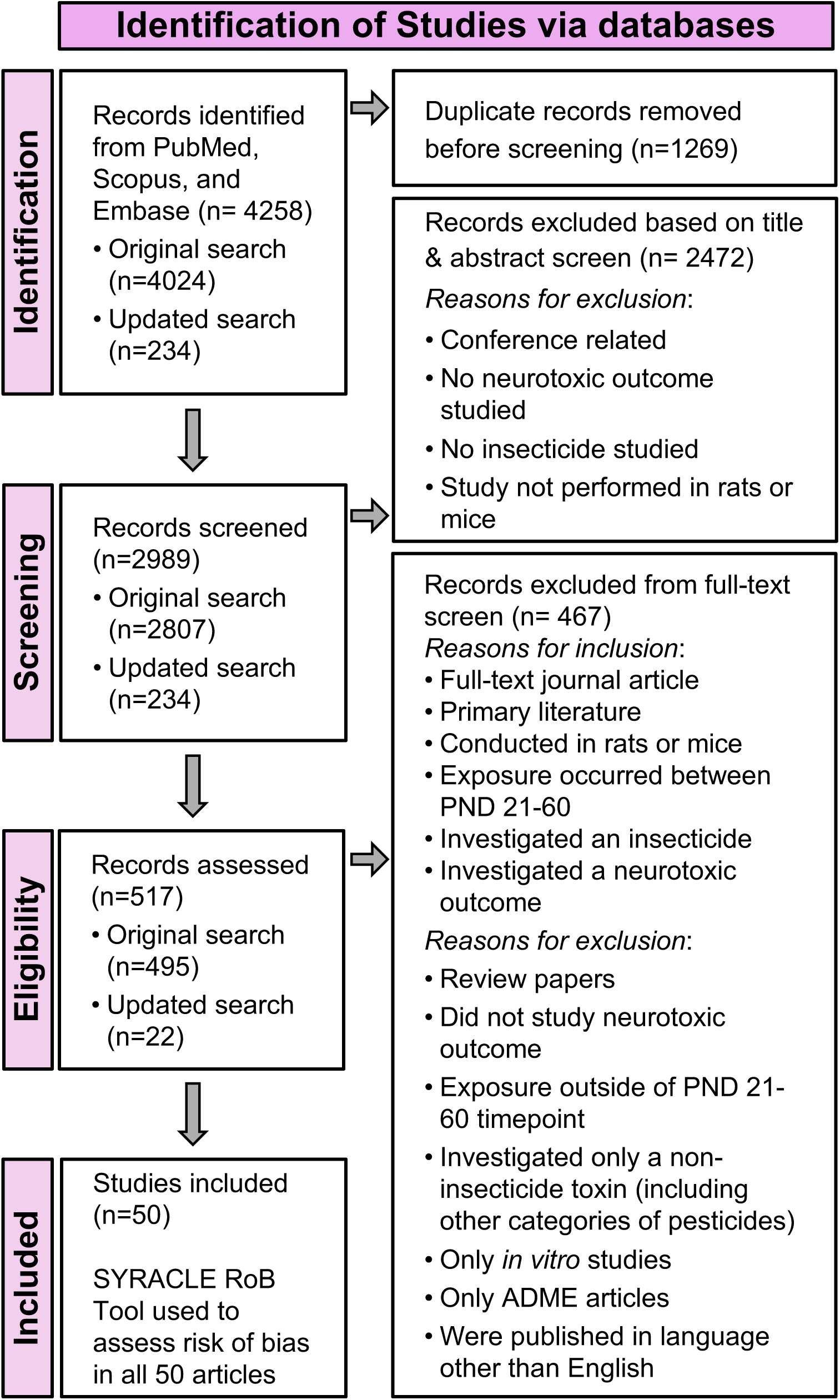
PRISMA flowchart of the screening strategy for articles returned from databases and number of studies excluded at each step.

Risk of bias assessment (Figure 3A) revealed most studies reported pre-specified outcomes (92%) and included all animals in their analysis (78%), but few provided details on randomization, housing, allocation concealment, or blinding (“unclear” ratings).

**Figure 3.**
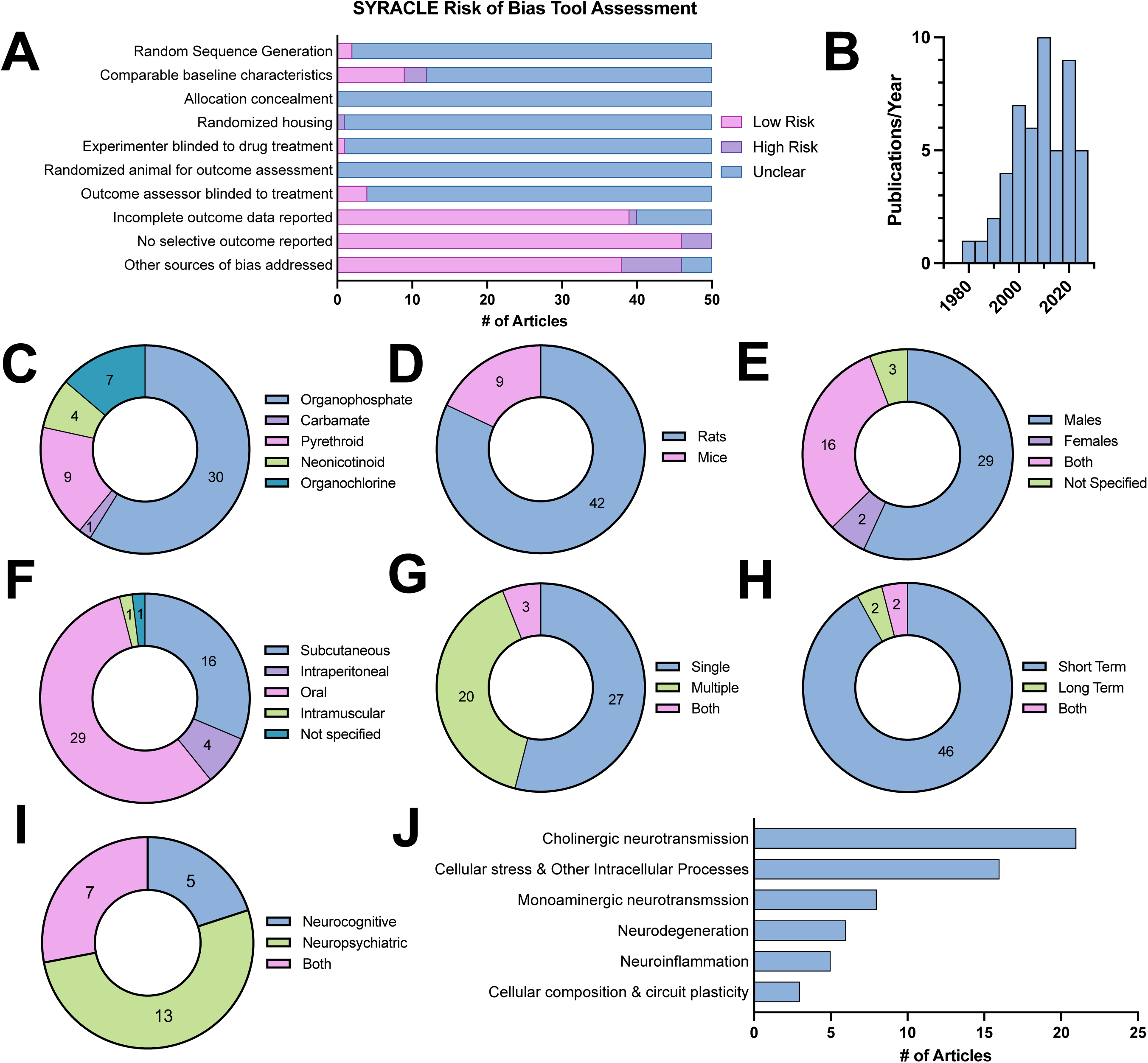
General characteristics of the studies included. **A)** Cumulative bar charts displaying risk of bias assessment across 10 items specified in the SYRCLE’s assessment tool. **B)** Histogram showing trend in articles published from 1980 to 2025 included in this review. Donut charts depicting the proportion and number of articles categorized by: **C)** Insecticide class (note that one article included study of two insecticides and was therefore double counted in this donut chart); **D)** Species; **E)** Sex; **F)** Administration route; **G)** Total number of doses; **H)** Outcome assessment timing (short term: during exposures and up to 1 week after final dose; long term > 1 week after final dose); **I)** Breakdown of articles assessing neurocognitive vs neuropsychiatric outcomes, among articles that investigated a neurobehavioral outcome; **J)** Breakdown of most common neurobiological outcomes measured.

To describe experimental diversity and scope, studies were categorized by insecticide class, species, sex, administration route, dosing frequency, and behavioral assessment timing (Figure 3C-I). Organophosphates were most frequently assessed (58.8% of studies), particularly chlorpyrifos (51.5% of organophosphate studies). Most studies used rats (84%), focused on males (58%, with 32% including both sexes), and employed oral (56.8%) or subcutaneous (31.4%) administration. Over half measured outcomes after a single dose (54%) and nearly all assessed short-term effects (92%).

Outcomes were grouped into four domains (see detailed definitions below): neurocognitive, neuropsychiatric, neurobiological, and general neurotoxicity. Many articles contributed to multiple domains. The most frequently reported neurobiological outcomes were cholinergic neurotransmission (21 articles), cellular stress and other intracellular processes (16), and monoaminergic neurotransmission (8, Figure 3J).

### Neurocognitive Outcomes

12 articles measured a neurocognitive outcome, which we defined as behavioral assays measuring learning and memory (Table 2). Learning and memory were assessed with Morris water maze, novel object recognition, Y-maze, or fear and avoidance conditioning tasks. Studies generally reported impairments in learning and memory across organophosphates, pyrethroids, and a single organochlorine (39–46). Neurocognitive performance was not impaired by neonicotinoid exposure (47, 48); however, there were only two studies.

**Table 2.** Summary of Neurocognitive Outcomes (learning and memory)

| ***Insecticide Class*** | ***Compounds Studied*** | ***# of Articles*** | ***Neurocognitive Outcomes*** | ***Consistency of Findings*** | ***References*** |
| --- | --- | --- | --- | --- | --- |
| Organophosphates | Chlorpyrifos, DFP, Dichlorvos, Malathion, Monocrotophos | 7 | Impairments in spatial learning, recognition memory, fear conditioning, and avoidance learning; some sex-specific or dose-dependent effects reported. | Impairments reported across multiple studies | (39-43, 55, 56) |
| Pyrethroids | Cypermethrin, Deltamethrin | 2 | Deficits in working memory and conditioned avoidance | Impairments reported across both studies | (44, 45) |
| Neonicotinoids | Clothianidin, Imidacloprid | 2 | No deficits in fear memory or working memory reported | No effects reported across both studies | (47, 48) |
| Organochlorines | Lindane | 1 | Impaired working memory and conditioned avoidance | Inconclusive (only a single study) | (46) |

### Neuropsychiatric Outcomes

21 articles assessed neuropsychiatric outcomes, defined as behavioral assays measuring anxiety- and depressive-relevant, locomotor, and other behaviors (Table 3). Locomotor activity was the most studied endpoint across insecticide classes, typically measured in an open field. Decreased activity was reported following exposure to organophosphates, carbamates, and a single organochlorine (46, 49–54). However, several studies observed no effect (55), including studies performed weeks following exposures (40, 54, 56). For a single pyrethroid studied, single doses had no effect (57), whereas repeated dosing increased locomotor activity (45, 58). Neonicotinoid exposure did not alter activity levels (47, 48). Collectively, these mixed findings suggest locomotor effects may depend on total exposure dose, insecticide class, and assessment timing.

**Table 3.**
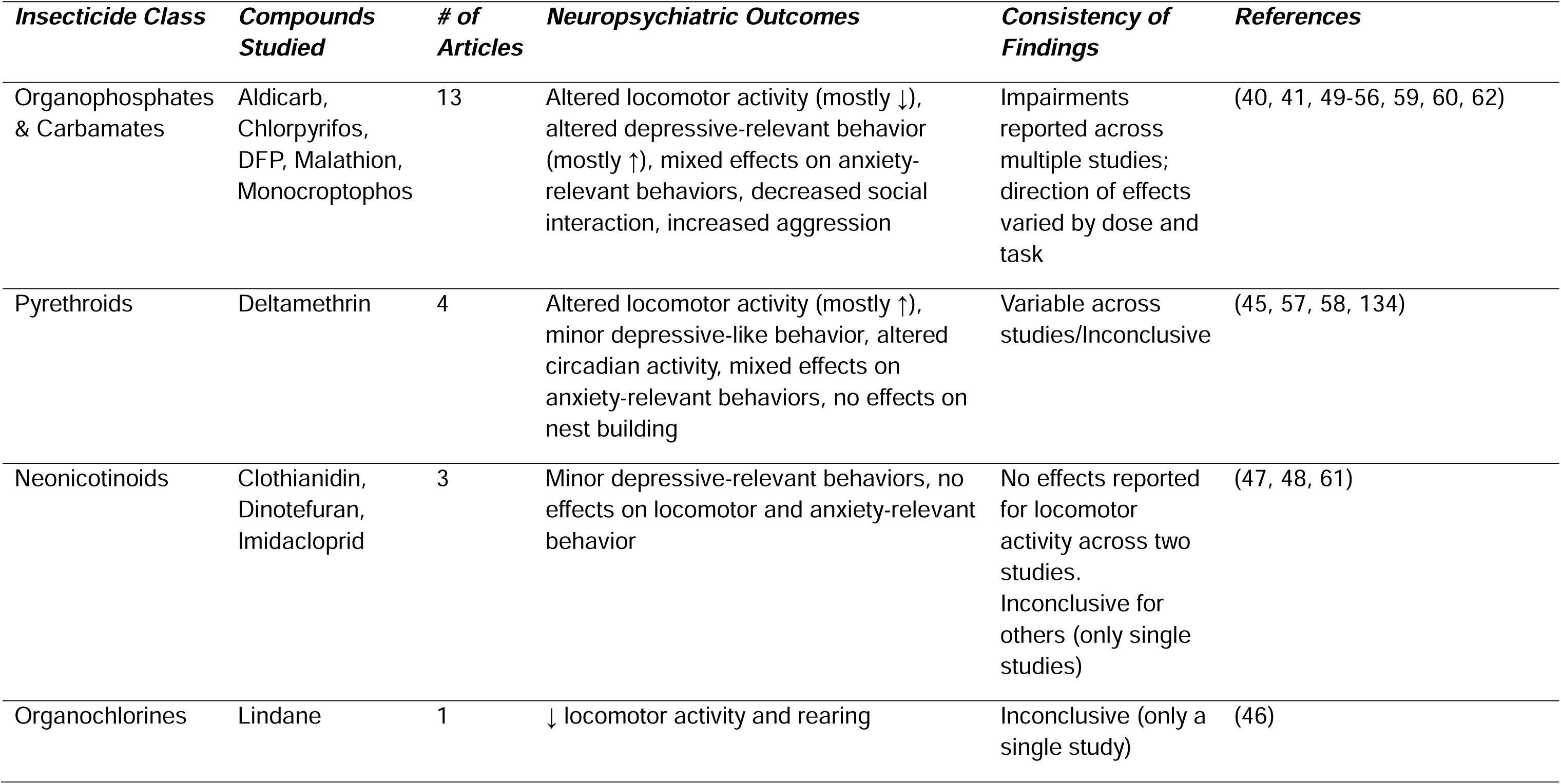
Summary of Neuropsychiatric Outcomes (anxiety- and depressive-relevant, locomotor, and other behaviors)

Depressive- and anxiety-relevant behaviors were assessed using forced swim, tail suspension, novelty suppressed feeding, elevated plus maze, or light/dark assay. Organophosphates and a single pyrethroid generally increased immobility on tail suspension or forced swim, though not consistently across doses (55, 58–60). In contrast, a neonicotinoid study reported decreased immobility (61). Findings for anxiety-relevant behavior were inconsistent. The assessment timing appeared to influence impacts of organophosphates: delayed assessment increased anxiety-relevant behavior, indicated by reduced time in the open arm of the elevated plus maze (41), whereas shorter-term assessment decreased anxiety-relevant behavior (50, 60). Other studies found no effect on anxiety-relevant measures (47, 55, 62). Importantly, performance indicators such as more open arm time in elevated plus maze or center time in open field are typically interpreted as reduced anxiety-relevant behavior. However, such outcomes may also capture impulsivity-relevant behavior (63), a distinction particularly relevant for insecticide exposure, given associated ADHD symptoms in humans (Table 1).

Single studies additionally showed adolescent insecticide exposure led to changes in other behaviors relevant to other neuropsychiatric disorders, such as in social interaction, aggression, and circadian rhythm (41, 58). No changes were observed in nest building following a pyrethroid exposure (58).

### Neurobiological Outcomes

40 articles measured neurobiological outcomes (Table 4). Most studied organophosphates and carbamates, reporting consistent decreases in acetylcholinesterase activity and alterations in cholinergic and monoaminergic signaling across brain regions. Monoaminergic systems including dopamine and serotonin were also affected across pyrethroids studies and organochlorines. Across insecticide classes, studies also reported evidence of oxidative stress (affected by all four classes), neuroinflammation via microglial or astrocyte changes (affected by organophosphates and carbamates, as well as pyrethroids), and neurodegenerative changes (with damaged and dying cells), indicating convergent disruption of neuronal and glial homeostasis. In a genome-wide assessment, a single organophosphate altered hippocampal synaptic plasticity and RNA splicing pathways among others (56).

**Table 4.** Summary of Neurobiological Outcomes (neurochemical, neuroinflammation, neurodegeneration, etc.)

| ***Insecticide Class*** | ***Compounds Studied*** | ***# of Articles*** | ***Neurobiological Outcomes*** | ***Consistency of Findings*** | ***References*** |
| --- | --- | --- | --- | --- | --- |
| Organophosphates & Carbamates | Aldicarb, Chlorpyrifos, DFP, Diazinon, Dichlorvos, Dichlofenthion, Fenitrothion, Malathion, Monocrotophos, Parathion, Trichlorfon, Clorfenvinphos | 26 | Altered cholinergic and monoaminergic signaling, oxidative stress, neurodegeneration, neuroinflammation, transcriptomic changes<br><u>Brain regions studied:</u> frontal cortex, striatum, hippocampus, thalamus, amygdala, temporal cortex, parietal cortex, piriform cortex, substantia nigra, medulla, cerebellum, spinal cord | Impairments reported across multiple studies | (39-43, 49, 50, 52-54, 56, 59, 70-73, 75, 84, 93, 94, 96, 135-139) |
| Pyrethroids | Cypermethrin, Deltamethrin | 5 | Altered monoamine systems, oxidative stress, neurodegeneration, neuroinflammation<br><u>Brain regions studied:</u> frontal cortex, striatum, hippocampus, pituitary, cerebellum, hypothalamus, medulla, hypothalamus | Impairments reported across multiple studies | (44, 45, 57, 58, 140) |
| Neonicotinoids | Dinotefuran, Imidacloprid, Clothianidin | 4 | Minor oxidative stress<br><u>Brain regions studied:</u> forebrain, hippocampus, dorsal raphe nucleus | Inconclusive (only single studies) | (47, 48, 61, 140) |
| Organochlorines | Chlordecone, DDT, Hepatochlor, Lindane | 6 | Altered monoaminergic neurotransmission, oxidative stress, neurodegeneration, gene expression & methylation changes<br><u>Brain regions studied:</u> frontal cortex, hippocampus, hypothalamus, thalamus, striatum, cerebellum, substantia nigra, medulla | Impairments reported across multiple studies | (46, 64, 65, 76, 95, 141) |

### General Neurotoxic Outcomes

18 articles measured a general neurotoxic outcome, which was defined here as seizures, tremors, and clinical signs (Table 5). Most articles studied organophosphates, carbamates, or pyrethroids and reported consistent occurrence of seizures, tremors, convulsions, altered neuromuscular function, and altered reflexes. Only four articles studied a neonicotinoid or an organochlorine and reported limited or short-lasting neurotoxic effects (47, 64–66). Importantly, a range of dosing was used, and neurotoxic symptoms typically occurred at higher dosages.

**Table 5.** Summary of General Neurotoxic Outcomes (seizures, tremors, neuromuscular function alterations)

| ***Insecticide Class*** | ***Compounds Studied*** | ***# of Articles*** | ***General Neurotoxic Outcomes*** | ***Consistency of Findings*** | ***References*** |
| --- | --- | --- | --- | --- | --- |
| Organophosphates & Carbamates | Aldicarb, Chlorpyrifos, DFP, Fenthion, Monocrotophos, Paraoxon | 10 | Seizures, tremors, gait/ataxia, impaired reflexes, altered neuromuscular function, other acute cholinergic signs | Impairments reported across multiple studies; severity varied by compound, dose, and treatment context | (39, 40, 43, 49, 51, 52, 54, 84, 111, 137) |
| Pyrethroids | Type I (Pyrethrins, Resmethrin, S-bioallethrin, Permethrin, Bifenthrin, Tefluthrin) & Type II (Cypermethrin, Esfenvalerate, β-Cyfluthrin, Deltamethrin, Fenpropathrin, λ-Cyhalothrin) | 4 | Tremors, salivation, convulsions, exaggerated reflexes, altered neuromuscular function | Impairments reported across multiple studies; both acute and dose-dependent effects observed | (57, 142-144) |
| Neonicotinoids | Clothianidin | 1 | Tremors observed after administration, short-lasting (<24h) | Inconclusive (only a single study) | (47) |
| Organochlorines | Lindane | 3 | Mixed findings: no tremors or seizures at moderate doses, but convulsions reported under some conditions | Variable findings across studies | (64-66) |

## 4. Discussion

### 4.1 Relevance of findings to mechanisms of psychiatric disorders

We focus here on discussing behavioral outcomes and neurobiological mechanisms of insecticide toxicity that may demonstrate increased psychiatric risk and provide recommendations for future studies.

#### Neurotransmitter Systems: Cholinergic and Monoaminergic Neurotransmission

Adolescence is a sensitive period for cholinergic and dopaminergic circuit maturation. Nicotinic acetylcholine receptor (nAChRs) levels peak during adolescence, particularly in striatum and midbrain, while choline acetyltransferase (ChAT), the key enzyme for acetylcholine synthesis, increases through adolescence in cortex and midbrain. Cholinergic projections modulate midbrain dopaminergic neurons and striatal neurons, undergoing substantial adolescent remodeling (8, 11). Maturation of these circuits supports cognitive and behavioral development and contributes to adolescent risk-taking and novelty-seeking (10, 67, 68). Disruption of these processes, such as with substance use, can induce short- and long-term memory, attention, and behavioral control impairments (69).

Rodent studies consistently report acetylcholinesterase inhibition in frontal cortex and striatum (70–73) paralleling peripheral acetylcholinesterase and butyrylcholinesterase activity inhibition in exposed humans (23, 25, 74). Insecticides also reduced ChAT activity and muscarinic receptor binding (43, 71, 73, 75) and alter monoaminergic signaling in dopaminergic, serotonergic, noradrenergic pathways in frontal cortex, striatum, and midbrain (53, 54, 58, 65, 76). *In vitro*, insecticides are directly toxic to dopaminergic neurons (53, 77). These findings collectively support insecticide-induced disruption of cholinergic and dopaminergic signaling critical for adolescent circuit maturation. Nicotine, which acts on nAChRs like neonicotinoids, induces persistent neurocircuitry changes linked to increased psychiatric risk (11). This and other insecticides, through enhancing neuronal firing or elevating synaptic acetylcholine, may similarly produce comparable long-term effects on brain function.

#### Inflammation & Neuroinflammation

Growing evidence implicates both central and peripheral immune systems in psychiatric disease. Elevated serum cytokine and chemokine levels have been reported in humans with mood disorders, schizophrenia, and cognitive impairment as well as mice exposed to chronic stress (78, 79). Insecticides appear to engage similar pathways: rodent studies showed adolescent pyrethroid exposure increases pro-inflammatory IL-1β, NF-κB, and IL-1R1 in cortex and hippocampus (44), and altered cytokine levels have been reported in humans exposed to insecticides (80, 81). Peripheral immune activation may also impact brain by compromising blood-brain-barrier (BBB) integrity, an outcome in mice exposed to chronic stress and with schizophrenia, depression, and autism spectrum disorder in humans (82, 83). Consistent with this, adolescent organophosphate exposure altered vasculature in cortex, striatum, and hippocampus consistent with compromised BBB permeability (84). Together, these findings suggest adolescent insecticide-induced immune activation and BBB disruption could have neuropsychiatric impacts.

Glia participate in immune responses and are critical regulators of myelination, synaptic structure and function, and neuronal migration which, in adolescence, particularly affects dopaminergic circuits in frontal cortex and striatum (85, 86). Shifting these processes out of their typical range can have lasting effects, as demonstrated by microglial depletion in prefrontal cortex during adolescence, but not adulthood, altering adult cognition (87, 88). Insecticide exposure may also disrupt these processes, as adolescent organophosphate exposure activates astrocytes and microglia in amygdala, hippocampus, and cortex (40, 41). Microglial activation also occurs in humans during major depression episodes (89), and reducing microglia-mediated neuroinflammation during adolescent chronic stress decreases neuronal damage and depressive-relevant behaviors in mice (90). Thus, adolescent insecticide-induced glial activation may disrupt key mesocorticolimbic circuit development, contributing to long-term cognitive and emotion regulation changes.

#### Cellular Stress

Oxidative stress has been extensively implicated in aging and neurodegenerative disease pathogenesis. More recently, together with neuroinflammation, its induction has been proposed as a contributor to psychiatric disease (91), with antioxidant treatment in adolescent mice protecting against risk (92). Oxidative stress is a well-studied insecticide mechanism and here, was a consistent outcome after adolescent exposure, including in frontal cortex, striatum, and hippocampus (46, 48, 50, 54, 56, 71, 93–96). These findings parallel increased oxidative stress serum measures in 6-12 year old children (97) or urinary measures in 8-11 year old children (98) exposed to insecticides as well as *in vitro* studies demonstrating oxidative stress in astrocytes, microglia, and neurons after insecticide exposure (53, 77, 99, 100).

#### Neurodegeneration & Cellular Composition Changes

Insecticide exposure has been linked to neurodegenerative disorders, particularly Parkinson’s disease (101, 102). While these disorders typically emerge later in life, animal studies suggest adolescent exposures may initiate neurodegenerative processes. For example, adolescent organophosphate exposure increased persistent fluoro-jade C staining of apoptotic, necrotic, and autophagic cells in amygdala and hippocampus (40), and chronic adolescent pyrethroid or organochlorine exposures induced amyloid-β, tau, and other Parkinson’s- and Alzheimer’s associated proteins in hippocampus and cortex (42, 44, 46). Whether these changes reflect early contributors to long-term neurodegenerative risk or more immediate psychiatric risk remains unclear, though more recent evidence suggests shared pathophysiology (103).

Adolescent exposures also lead to changes in cellular composition, including reduced cortical and hippocampal parvalbumin-expression interneurons (PV-INs) and decreased hippocampal neuron counts and plasticity markers (41, 56). PV-INs regulate cortical activity and are critical for attention and cognition (104). Because PV-INs develop throughout adolescence, their disruption in rodent adolescent prefrontal cortex impairs hippocampal input integration and leads to adult cognitive deficits (105). Furthermore, PV-IN dysfunction in prefrontal cortex and hippocampus has been broadly implicated in psychiatric disorders (106), suggesting insecticide-induced changes during adolescence may contribute to long-term risk.

### 4.2 Limitations in Reviewed Studies and Considerations for Future Studies

#### Sex differences

Less than half of reviewed articles included both males and females. While many insecticide-induced impairments were shared across sexes, some were sex-specific. Males showed more pronounced deficits in fear learning (40) and greater seizure severity (40), whereas females showed effects at lower doses, including heightened reflexes (51) and delayed learning following neonicotinoid exposure (47). Organophosphate-induced neuroinflammation and neurodegeneration were observed in both sexes but appeared more pronounced in males (40). Similarly, organochlorine exposure increased dopamine transporter binding in males shortly after exposure, with effects emerging later across sexes (76). Collectively, these findings suggest male vulnerability to acute, severe outcomes, and female sensitivity to lower doses and delayed effects. However, methodological variability limits the ability to draw broad conclusions despite the importance of sex differences in neurodevelopmental and neuropsychiatric disorders and typical adolescent neurodevelopment (8, 11, 107–109). In human studies, comparable insecticide exposure levels have been reported in boys and girls (22, 74, 110), suggesting animal studies should examine both sexes and can appropriately model real-world exposures with similar dosages across males and females.

#### Exposure Methodology

We found a wide variety of insecticides studied and exposure methods (dosage, administration routes). This variability reflects complexities of studying environmental toxicants, as insecticides, even within a class, have differing levels of toxicity. Most articles reviewed here selected their study dosage based on lethal dose. For example, many rodent studies measure outcomes after a single, high dose calculated as percent of lethal dose (47, 70, 71, 111). These are relevant for acute poisonings in humans but do not reflect real-world exposures for most adolescents, who experience lower, more chronic exposures (13) which may have more nuanced psychiatric impacts. Studies of specific insecticides at specific doses have relevance to different populations, since those adolescents occupationally exposed to or living near agricultural areas have higher exposures (110, 112), and rural and urban children are mainly exposed to different insecticides (110, 113). Regardless, insecticide exposures are ubiquitous, and future studies must take modeling real-world exposure levels and routes into account.

#### Selecting Neurobehavioral and Neurobiological Measures

Selecting and appropriately interpreting behavioral outcomes can be difficult in animal models, and especially so in modeling depression and anxiety risk. In reviewed studies, researchers used tasks such as forced swim and tail suspension which have come under criticism in recent years for having limited validity (114). In the case of insecticides with clear motor/neuromuscular impacts, it becomes even more imperative to cautiously interpret movement-based assays. It will be important for future studies to assess and interpret behavior cautiously across multiple paradigms. To enhance translational relevance, rodent tasks relevant to observations in human populations, such as cognitive impairments and ADHD-relevant behaviors, using tests of learning, working memory, impulsivity, or behavioral inhibition, are appropriate. As discussed previously, paradigms traditionally used to measure anxiety-relevant behaviors may also have relevance to impulsivity-relevant behaviors. However, epidemiological research on adolescent insecticide exposure is still limited, and few studies have examined depressive-relevant outcomes despite consistent findings in rodent studies. Thus, rodent findings can also help guide future epidemiological research priorities.

We identified limited studies using -omics techniques to measure neurobiological outcomes. There were three studies, investigating: hypothalamic methylation changes (95), hippocampal transcriptomics (56), and proteomics (44). Such techniques offer unbiased insight into and hypothesis generation about toxicity mechanisms as well as potential for translation to human studies, in which peripheral DNA methylation changes have been associated with high insecticide exposure (115). Additionally, we identified limited circuit-level studies which may have more direct translation to growing appreciation of circuit-level dysfunction in neuropsychiatric illness (116); as neuroscience tools advance, techniques like *in vivo* electrophysiology and fiber photometry can shed light on how insecticides impact neural circuits in real time.

#### Exposure and Assessment Age & Limitations

It is possible our search strategy did not identify all relevant studies. This limitation also reflects the greater challenge of defining rodent “adolescence.” Researchers may describe rodents prior to PD60 as “adults,” while the brain is still undergoing neurodevelopmental processes similar to those in human adolescence (117). We were not able to draw conclusions about more specific exposure timing during adolescence, but this will be relevant as studies provide more evidence in the future, since toxicity may impact differing neurodevelopmental processes depending on when they are occurring (Figure 1). Furthermore, long-term sequelae of adolescent insecticide exposure remain unclear, although impacts within adolescence itself is an important one for the health of young people. Overall, greater clarity of cross-species neurodevelopmental stages could allow for more accurate interpretation and implications of findings in rodent models.

### 4.3 Knowledge gaps & need for modeling real-world exposures

Several gaps require more research to elucidate insecticide impacts on adolescent psychiatric vulnerability. One central research need is modeling real-world exposures to allow for translational insight. Insecticide exposure levels below regulatory agency guidelines may still contribute to neurobehavioral outcomes over time. To address this, researchers should use human exposure measurements when designing exposure methodology for animal studies.

Human exposure levels have been estimated through urinary insecticide metabolites, acetylcholinesterase activity, or insecticide detection on wearable wristbands (21, 22, 74), giving basic researchers a prime opportunity to model translational exposures in animals (118).

Globally, longitudinal child and adolescent study cohorts examine agrochemical exposure effects, including neurobehavioral outcomes. These include, but are not limited, to: ESPINA study in Ecuador (119), Salud Ambiental Montevideo study in Uruguay (120), Hired Child Farmworker Study in the United States (121), and Menoufia Governorate study in Egypt (122). Longitudinal studies are opportunities for transdisciplinary collaboration to better understand psychiatric risk by engaging mental health epidemiologists, neuroscientists, and toxicologists (123). Particularly, studying newer insecticides, such as neonicotinoids, is needed both epidemiologically and in animal models. Although these compounds were developed to be less toxic, the current lack of evidence should not be interpreted as conclusive safety.

Another key research need is incorporation of the exposome framework, which is defined as totality of exposure individuals experience over their lives and how these combined exposures affect health (124). Studies here examined single insecticide effects, but real-world exposures consist of combined environmental factors—at the physical-chemical level (e.g., extreme temperatures, pesticides), ecosystem level (e.g., population density), lifestyle level (e.g., diet, drug use), and social level (e.g., psychosocial stress). Such combinations overlaid on already complex genetic diversity makes studying psychiatric disease difficult but remains critical to understanding brain disorders. This exposome framework has begun to be applied within psychiatric research (125, 126). Unlike many stressors with unclear neural mechanisms, insecticides act through well-defined targets, suggesting they are powerful tools for probing how specific neurobiological disruptions contribute to psychiatric outcomes.

## 5. Conclusion

Adolescence is a unique vulnerability period for environmental exposures, highlighting a critical need to understand how ubiquitous environmental contaminant exposures may disrupt normal brain development and contribute to long-term psychiatric outcomes. Here, we performed a systematic review on neurobehavioral and neurobiological effects of adolescence insecticide exposure, demonstrating psychiatric-relevant outcomes convergent with reported human outcomes. Moreover, several identified insecticide-induced neurotoxicity mechanisms may disrupt adolescent brain development. While both epidemiological and animal studies on insecticides have historically focused on early development, we propose adolescence as another critical neurodevelopmental window and urge more research on this stage of psychiatric risk. Advancing our understanding of mechanisms of environmental exposures may provide new insights into modifiable risk factors contributing to the current youth mental health crisis. With such insight, we can design better prevention and mitigation strategies to minimize the burden of mental health disorders.

## Acknowledgements

We thank Riley Samuelson, University of Iowa Hardin Library for the Health Sciences, for help with the systematic search. Figures 1 and 4 were created using BioRender. This work was supported by the National Institutes of Health (NIEHS R01 ES035696 [all authors], NIEHS P30 ES005605 [all authors], NIH T32-NS007421 [MXC]).

**Figure 4.**
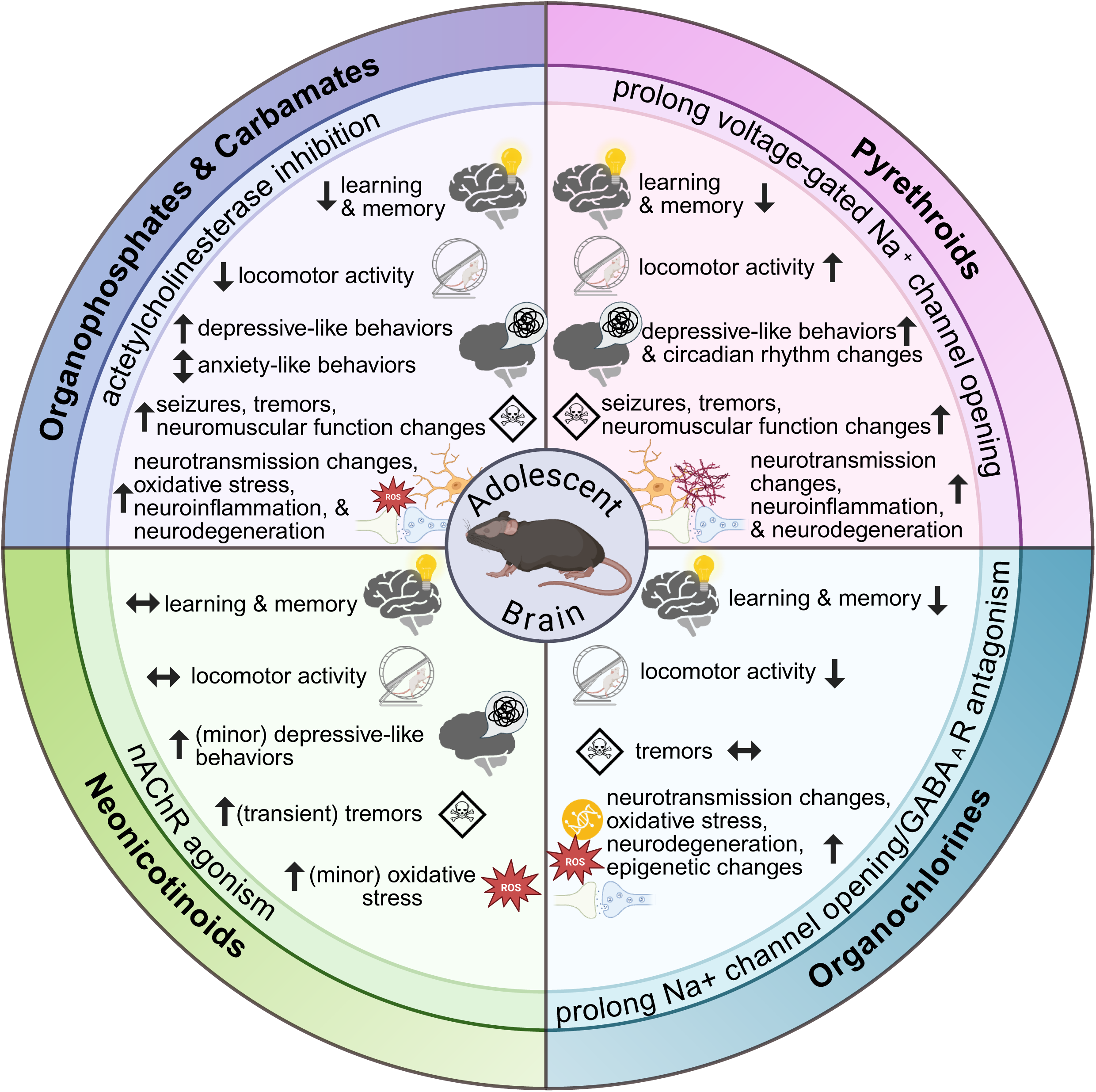
Cross-domain summary of neurotoxic effects from adolescent insecticide exposure in rodent models. *Made with BioRender*.

