## Supplementary Materials for "Psychiatric risk implications of adolescent exposure to environmental insecticides: a systematic review of rodent studies"

***Psychiatric risk implications of adolescent exposure to environmental insecticides: a systematic review of rodent students SEARCH TERMS***

**Pubmed**

**“Murinae”[Mesh]** OR “Rodent*”[Title/Abstract] OR “Pup”[Title/Abstract] OR “pup (rodent)”[Title/Abstract] OR “Rat”[Title/Abstract] OR “Rats”[Title/Abstract] OR “Murinae”[Title/Abstract] OR “Rattus”[Title/Abstract] OR “Mus”[Title/Abstract] OR “mus musculus”[Title/Abstract] OR “Mouse*”[Title/Abstract] OR “Mice”[Title/Abstract] OR “rattus rattus”[Title/Abstract] OR “murine”[Title/Abstract] OR “in vivo”[Title/Abstract]

AND

**“Adolescent”[Mesh]** OR “Adolescen*”[Title/Abstract] OR “Juvenile*”[Title/Abstract] OR “immature”[Title/Abstract] OR “young”[Title/Abstract]

AND

**“Pesticides”[Mesh:NoExp] OR “Insecticide*”[Mesh]** OR “Pesticide*”[Title/Abstract] OR “Insecticide*”[Title/Abstract] OR “Organophosphate*”[Title/Abstract] OR “Organophosphorus”[Title/Abstract] OR “Organic pesticide*”[Title/Abstract] OR “Organic phosphate*”[Title/Abstract] OR “phosphorus insecticide*”[Title/Abstract] OR “Organosulfur*”[Title/Abstract] OR “Carbamate*”[Title/Abstract] OR “Nicotinoid*”[Title/Abstract] OR “neonicotinoid*”[Title/Abstract] OR “Pyrethroid*”[Title/Abstract] OR “Organochlorine*”[Title/Abstract] OR “Chlorinated hydrocarbon*”[Title/Abstract] OR “Chlorinated organic*”[Title/Abstract] OR “chlorinated insecticide*” OR “chlorinated synthetic*”[Title/Abstract] OR “Formamidine*”[Title/Abstract] OR “Agrichemical*”[Title/Abstract] OR “Agrochemical*”[Title/Abstract] OR “Agricultural Chemical*”[Title/Abstract] OR “Chemical, Agricultural”[Title/Abstract] OR “Chemicals, Agricultural”[Title/Abstract]

AND

**“Mental Disorders”[Mesh]** OR **“Growth and Development”[Mesh]** OR **“Nervous System”[Mesh]** OR “Neurodevelopment”[Title/Abstract] OR “Neurodeve*”[Title/Abstract] OR “Neurobehavior”[Title/Abstract] OR “Neurobehaviour”[Title/Abstract] OR “Neurobehav*”[Title/Abstract] OR “Neurobehavioral”[Title/Abstract] OR “Neurobehavioural”[Title/Abstract] OR “Attention Deficit Disorder with Hyperactivity”[Title/Abstract] OR “Attention Deficit Disorder”[Title/Abstract] OR “ADHD”[Title/Abstract] OR “ADHD-like”[Title/Abstract] OR “Hyperactivity disorder”[Title/Abstract] OR “Deficit hyperactiv*”[Title/Abstract] OR “ADD”[Title/Abstract] OR “Hyperkinetic disorder”[Title/Abstract] OR “Hyperkinetic syndrome”[Title/Abstract] OR “attention deficit hyperactivity disorder”[Title/Abstract] OR “Attention deficit”[Title/Abstract] OR “Attention-deficit/hyperactivity disorder”[Title/Abstract] OR “Attention-deficit hyperactivity disorder”[Title/Abstract] OR “Toxic”[Title/Abstract] OR “Toxi*”[Title/Abstract] OR “neurodevelopmental disorder*”[Title/Abstract] OR “Neurotox*”[Title/Abstract] OR “Neurocognitive Disorder*”[Title/Abstract] OR “Mental disease*”[Title/Abstract] OR “Neurocogn*” [Title/Abstract] OR “Growth and Development”[Title/Abstract] OR “Growth”[Title/Abstract] OR “Develop*”[Title/Abstract] OR “Brain*”[Title/Abstract] OR “Central Nervous System”[Title/Abstract] OR “CNS”[Title/Abstract]

OR “Nervous System”[Title/Abstract] OR “growth, development and aging”[Title/Abstract] OR “psychiatric”[Title/Abstract] OR “psychological”[Title/Abstract] OR “neuropsychiatric”[Title/Abstract] OR “Mood disorder*”[Title/Abstract] OR “depressive*”[Title/Abstract] OR “depression*”[Title/Abstract] OR “Substance-Related Disorder*”[Title/Abstract] OR “Substance use disorder*”[Title/Abstract] OR “addict*”[Title/Abstract] OR “Alcohol use disorder”[Title/Abstract] OR “Alcohol-related disorder*”[Title/Abstract] OR “alcohol-self administration”[Title/Abstract] OR “alcohol-seeking”[Title/Abstract] OR “nicotine self-administration”[Title/Abstract] OR “nicotine seeking”[Title/Abstract] OR “tobacco use disorder”[Title/Abstract] OR “Cocaine-related disorder”[Title/Abstract] OR “Cocaine-self administration”[Title/Abstract] OR “cocaine-seeking”[Title/Abstract] OR “schizophrenia*”[Title/Abstract] OR “anxiety*”[Title/Abstract] OR “anxio*”

**Embase**

**‘rodent’/exp** OR ‘Rodent*’:ab,ti,kw OR ‘Pup’:ab,ti,kw OR ‘pup (rodent)’:ab,ti,kw OR ‘Rat’:ab,ti,kw OR ‘Rats’:ab,ti,kw OR ‘Murinae’:ab,ti,kw OR ‘Rattus’:ab,ti,kw OR ‘Mus’:ab,ti,kw OR ‘mus musculus’:ab,ti,kw OR ‘Mouse*’:ab,ti,kw OR ‘Mice’:ab,ti,kw OR ‘rattus rattus’:ab,ti,kw OR ‘murine’:ab,ti,kw OR ‘in vivo’:ab,ti,kw

AND

**‘juvenile’/exp** OR ‘Adolescen*’:ab,ti,kw OR ‘Juvenile*’:ab,ti,kw OR ‘immature’:ab,ti,kw OR ‘young’:ab,ti,kw

AND

**‘pesticide’/de** OR **‘insecticide’/exp** OR ‘Pesticide*’:ab,ti,kw OR ‘Insecticide*’:ab,ti,kw OR ‘Organophosphate*’:ab,ti,kw OR ‘Organophosphorus’:ab,ti,kw OR ‘Organic pesticide*’:ab,ti,kw OR ‘Organic phosphate*’:ab,ti,kw OR ‘phosphorus insecticide*’:ab,ti,kw OR ‘Organosulfur*’:ab,ti,kw OR ‘Carbamate*’:ab,ti,kw OR ‘Nicotinoid*’:ab,ti,kw OR ‘neonicotinoid*’:ab,ti,kw OR ‘Pyrethroid*’:ab,ti,kw OR ‘Organochlorine*’:ab,ti,kw OR ‘Chlorinated hydrocarbon*’:ab,ti,kw OR ‘Chlorinated organic*’:ab,ti,kw OR ‘chlorinated insecticide*’ OR ‘chlorinated synthetic*’:ab,ti,kw OR ‘Formamidine*’:ab,ti,kw OR ‘Agrichemical*’:ab,ti,kw OR ‘Agrochemical*’:ab,ti,kw OR ‘Agricultural Chemical*’:ab,ti,kw OR ‘Chemical, Agricultural’:ab,ti,kw OR ‘Chemicals, Agricultural’:ab,ti,kw

AND

**‘mental disease’/exp** OR **‘toxicity’/exp** OR **‘growth, development and aging’/exp** OR **‘nervous system’/exp** OR ‘Neurodevelopment’:ab,ti,kw OR ‘Neurodeve*’:ab,ti,kw OR ‘Neurobehavior’:ab,ti,kw OR ‘Neurobehaviour’:ab,ti,kw OR ‘Neurobehav*’:ab,ti,kw OR ‘Neurobehavioral’:ab,ti,kw OR ‘Neurobehavioural’:ab,ti,kw OR ‘Attention Deficit Disorder with Hyperactivity’:ab,ti,kw OR ‘Attention Deficit Disorder’:ab,ti,kw OR ‘ADHD’:ab,ti,kw OR ‘ADHD-like’:ab,ti,kw OR ‘Hyperactivity disorder’:ab,ti,kw OR ‘Deficit hyperactiv*’:ab,ti,kw OR ‘ADD’:ab,ti,kw OR ‘Hyperkinetic disorder’:ab,ti,kw OR ‘Hyperkinetic syndrome’:ab,ti,kw OR ‘attention deficit hyperactivity disorder’:ab,ti,kw OR ‘Attention deficit’:ab,ti,kw OR ‘Attention-deficit/hyperactivity disorder’:ab,ti,kw OR ‘Attention-deficit hyperactivity disorder’:ab,ti,kw OR ‘Toxic’:ab,ti,kw OR ‘Toxi*’:ab,ti,kw OR ‘neurodevelopmental disorder*’:ab,ti,kw OR ‘Neurotox*’:ab,ti,kw OR ‘Neurocognitive Disorder*’:ab,ti,kw OR ‘Mental disease*’:ab,ti,kw OR ‘Neurocogn*’:ab,ti,kw OR ‘Growth and Development’:ab,ti,kw OR ‘Growth’:ab,ti,kw OR ‘Develop*’:ab,ti,kw OR ‘Brain*’:ab,ti,kw OR ‘Central Nervous System’:ab,ti,kw OR ‘CNS’:ab,ti,kw OR ‘Nervous System’:ab,ti,kw OR ‘growth, development and aging’:ab,ti,kw OR ‘psychiatric’:ab,ti,kw OR ‘psychological’:ab,ti,kw OR ‘neuropsychiatric’:ab,ti,kw OR ‘Mood disorder*’:ab,ti,kw OR ‘depressive*’:ab,ti,kw OR ‘depression*’:ab,ti,kw OR ‘Substance-Related Disorder*’:ab,ti,kw OR ‘Substance use disorder*’:ab,ti,kw OR ‘addict*’:ab,ti,kw OR ‘Alcohol use disorder’:ab,ti,kw OR ‘Alcohol-related disorder’:ab,ti,kw OR ‘alcohol-self administration’:ab,ti,kw OR ‘alcohol-seeking’:ab,ti,kw OR ‘nicotine self-administration’:ab,ti,kw OR ‘nicotine seeking’:ab,ti,kw OR ‘tobacco use disorder’:ab,ti,kw OR ‘Cocaine-related disorder’:ab,ti,kw OR ‘Cocaine-self administration’:ab,ti,kw OR ‘cocaine-seeking’:ab,ti,kw OR ‘schizophrenia*’:ab,ti,kw OR ‘anxiety*’:ab,ti,kw OR “anxio*”:ab,ti,kw

**SCOPUS**

In Title, Abstract, Keyword search

“Rodent*” OR “Pup” OR “pup (rodent)” OR “Rat” OR “Rats” OR “Murinae” OR “Rattus” OR “Mus” OR “mus musculus” OR “Mouse*” OR “Mice” OR “rattus rattus” OR “murine” OR “in vivo”

**AND** “Adolescen*” OR “Juvenile*” OR “immature” OR “young”

**AND** “Pesticide*” OR “Insecticide*” OR “Organophosphate*” OR “Organophosphorus” OR “Organic pesticide*” OR “Organic phosphate*” OR “phosphorus insecticide*” OR “Organosulfur*” OR “Carbamate*” OR “Nicotinoid*” OR “neonicotinoid*” OR “Pyrethroid*” OR “Organochlorine*” OR “Chlorinated hydrocarbon*” OR “Chlorinated organic*” OR “chlorinated insecticide*” OR “chlorinated synthetic*” OR “Formamidine*” OR “Agrichemical*” OR “Agrochemical*” OR “Agricultural Chemical*” OR “Chemical, Agricultural” OR “Chemicals, Agricultural”

**AND** “Neurodevelopment” OR “Neurodeve*” OR “Neurobehavior” OR “Neurobehaviour” OR “Neurobehav*” OR “Neurobehavioral” OR “Neurobehavioural” OR “Attention Deficit Disorder with Hyperactivity” OR “Attention Deficit Disorder” OR “ADHD” OR “ADHD-like” OR “Hyperactivity disorder” OR “Deficit hyperactiv*” OR “ADD” OR “Hyperkinetic disorder” OR “Hyperkinetic syndrome” OR “attention deficit hyperactivity disorder” OR “Attention deficit” OR “Attention-deficit/hyperactivity disorder” OR “Attention-deficit hyperactivity disorder” OR “Toxic” OR “Toxi*” OR “neurodevelopmental disorder*” OR “Neurotox*” OR “Neurocognitive Disorder*” OR “Mental disease*” OR “Neurocogn*” OR “Growth and Development” OR “Growth” OR “Develop*” OR “Brain*” OR “Central Nervous System” OR “CNS” OR “Nervous System” OR “growth, development and aging” OR “psychiatric” OR “psychological” OR “neuropsychiatric” OR “Mood disorder*” OR “depressive*” OR “depression*” OR “Substance-Related Disorder*” OR “Substance use disorder*” OR “addict*” OR “Alcohol use disorder” OR “Alcohol-related disorder” OR “alcohol-self administration” OR “alcohol-seeking” OR “nicotine self-administration” OR “nicotine seeking” OR “tobacco use disorder*” OR “Cocaine-related disorder*” OR “Cocaine-self administration” OR “cocaine-seeking” OR “schizophrenia*” OR “anxiety*” OR “anxio*”

**Table S1.** Neurocognitive Outcomes (learning and memory) & Methods

| **Insecticide** | **Animal Model** | **Route & Dose** | **Exposure Age** | **Assessment Age** | **Effects on Behavior** | **Reference** |
| --- | --- | --- | --- | --- | --- | --- |
| ***Organophosphates*** | | | | | | |
| Chlorpyrifos | Rat, Long Evans,  Both sexes  n=7-8/group | Vehicle vs 0.3 vs 7 mg/kg for two doses  Subcutaneous | PD 22 and 26 | PD 24-28 | **MORRIS WATER MAZE:**  ↑ escape latency, all doses | (1) |
|  | Rat, Sprague Dawley, Male  n=12/group | Vehicle vs 2.5 or 5 or 10 or 20 mg/kg/day for 10 days  Subcutaneous | PD 27-36 | PD 37-52 | **CONDITIONED AVOIDANCE:**  ↑ escape failures at 5, 10, and 20 mg/kg/day; ↔ at 2.5 mg/kg/day | (2) |
| DFP | Rat, Sprague Dawley, Both sexes  n= 10-12/sex/group | Vehicle vs 3.4 mg/kg for a single dose  Subcutaneous  *Post-treatment: atropine sulfate and pralidoxime* | PD 28 | ~ PD 56 | **NOVEL OBJECT RECOGNITION:**  ↓ discrimination index, both sexes  **CONTEXTUAL & CUED FEAR CONDITIONING:**  ↑ fear-associated memory deficits, both sexes, more pronounced in males | (3) |
|  | Rat, Sprague Dawley, Males  n=8-13/group | Vehicle vs 1.4 x LD50 mg/kg  Subcutaneous  *Pre-treatment: pyridostigmine bromide;*  *Post-treatment: 2-PAM and atropine methylnitrate* | PD 28 | ~ 7 & 14 months | **NOVEL OBJECT RECOGNITION:**  ↓ time with novel object at 7 and 14 months  **MORRIS WATER-MAZE:**  ↓ latency to platform (timepoint of testing not specified) | (4) |
| Dichlorvos | Rat, Sprague Dawley, Male  n=8-10/group | Vehicle vs 0.1 vs. 2 mg/kg/day for 28 days  Subcutaneous | PD 21-48 | Not specified; presumably PD 49-55 | **MORRIS WATER MAZE:**  ↓ escape latency at 0.1 mg/kg/day; ↔ at 2 mg/kg/day | (5) |
| Malathion | Mouse, C57BL6, Males  n=9/group | Vehicle vs 50 mg/kg/day for 14 days  Oral (gavage) | 4-5 weeks | 6-7 weeks | **NOVEL OBJECT RECOGNITION:**  ↔ discrimination and recognition index | (6) |
| Monocrotophos | Rat, Wistar, Female  n=4-5/group | Vehicle vs 0.5 or 1 mg/kg/day for 27 days  Oral (gavage) | PD 22-49 | PD 50 | **CONDITIONED AVOIDANCE:**  ↓ avoidance behavior, all doses | (7) |
|  | Rat, Wistar, Female  n=4-5/group | Vehicle vs 0.5 or 1 mg/kg/day for 27 days  Oral (gavage) | PD 22-49 | PD 65 | **CONDITIONED AVOIDANCE:**  ↓ avoidance behavior, all doses | (7) |
| ***Pyrethroids*** | | | | | | |
| Cypermethrin | Rat, Wistar, Males  n=20/group | Vehicle vs 10 or 25 mg/kg/day for 2 weeks  Oral (gavage) | PD 24 | PD 38-45 | **Y-MAZE:**  ↑ errors during learning phase, all doses  ↓ % saving memory up to 7 days post learning, all doses  **CONDITIONED AVOIDANCE:**  ↓ learning in all retention trials, all doses | (8) |
| Deltamethrin | Rat, Wistar, Male  n=10/group | Vehicle vs 7 mg/kg/day for 15 days  Oral (gavage) | PD 22-37 | PD 38-40 | **CONDITIONED AVOIDANCE:**  ↓ % avoidance response, all doses | (9) |
| ***Neonicotinoids*** | | | | | | |
| Clothianidin | Mouse, C57Bl/6N, Both sexes  n=7-10/group | Vehicle vs 80 mg/kg for a single dose  Oral (gavage) | 6 weeks | 3 months | **CONTEXTUAL/CUED FEAR CONDITIONING:**  ↔ conditional freezing, both sexes  ↔ contextual freezing, both sexes  ↔ cued freezing, both sexes | (10) |
|  | Mouse, C57Bl/6N, Both sexes  n=6-10/group | Vehicle vs 80 mg/kg for a single dose  Oral (gavage) | 6 weeks | 7 months | **CONTEXTUAL/CUED FEAR CONDITIONING:**  ↔ conditional freezing, both sexes  ↔ contextual freezing in males; ↓ in females  ↔ cued freezing, both sexes | (10) |
| Imidacloprid | Mouse, Swiss albino, Male  n=6/group | Vehicle vs 1 or 2 mg/kg/day for 40 days  Oral (gavage) | PD 21-60 | PD 56-60 | **MORRIS WATER MAZE:**  ↔ latency to find platform, both sexes | (11) |
| ***Organochlorine*** | | | | | | |
| Lindane | Rat, Wistar, Male  n=6/group | Vehicle vs 2.5 mg/kg/day for 21 days  Oral | 3 weeks | Not specified; presumably 6 weeks | **Y-MAZE:**  ↓ % alternations  **CONDITIONED AVOIDANCE:**  ↓ % conditioned avoidance response | (12) |

**Table S2.** Neuropsychiatric outcomes (anxiety- and depressive-relevant, locomotor, and other behaviors) & Methods

| **Insecticide** | **Animal Model** | **Route & Dose** | **Exposure Age** | **Assessment Age** | **Effects on Behavior** | **Reference** |
| --- | --- | --- | --- | --- | --- | --- |
| ***Organophosphates & Carbamates*** | | | | | | |
| Aldicarb | Rat, Long Evans, Both sexes  n=10/group | Vehicle vs 0.08 or 0.15 or 0.26 mg/kg for a single dose  Oral (gavage) | PD 27 | PD 27 & 28 | **OPEN FIELD TEST** (as part of Functional Observational Battery), 1 HR following exposure:  ↓ locomotion at 0.15 and 0.26, both sexes  ↓ rearing at 0.15 and 0.26, both sexes  **OPEN FIELD TEST** (as part of Functional Observational Battery), 24 HR following exposure:  ↔ locomotion or rearing, all doses, both sexes | (13) |
| Chlorpyrifos | Rat, Sprague Dawley, Male  n=8/group | Vehicle vs 2.5 or 5 or 10 mg/kg/day for 10 days  Subcutaneous | PD 22-32 | PD 33 | **FORCED SWIM TEST:**  ↔ immobility time at 2.5 and 5 mg/kg/day; ↑ at 10 mg/kg/day | (14) |
|  | Rat, Sprague Dawley, Male  n=10/group | Vehicle vs 7.45 mg/kg/day for 37 days  Intraperitoneal | PD 23-60 | PD 59-60 | **OPEN FIELD TEST:**  ↓ locomotion  **FORCED SWIM TEST:**  ↓ immobility time; ↑ swimming time  **ELEVATED PLUS MAZE:**  ↑ total entries | (15) |
|  | Rat, Sprague Dawley, Male  n=12/group | Vehicle vs 2.5 or 5 or 10 or 20 mg/kg/day for 10 days  Subcutaneous | PD 27-36 | PD 37-52 | **FORCED SWIM TEST:**  ↑ immobility time at 10 mg/kg/day; ↔ at 2.5, 5, and 20 mg/kg/day  **OPEN FIELD TEST:**  ↔ locomotion, all doses  **NOVELTY SUPPRESSED FEEDING:**  ↔ feeding latency period, all doses | (2) |
|  | Rat, Sprague Dawley, Male  n=9-10/group | Vehicle vs 10 or 20 or 40 or 80 or 160 mg/kg/day for 7 days  Subcutaneous | PD 29-35 | PD 37-46 | **FORCED SWIM TEST:**  ↑ immobility time at 10 mg/kg/day; ↔ at 20, 40, and 80 mg/kg/day; ↓ at 160 mg/kg/day  **ELEVATED PLUS MAZE:**  ↑ open arm time at 40, 80, and 160 mg/kg; ↔ at 10 and 20 mg/kg  **NOVELTY SUPPRESSED FEEDING:**  ↓ latency period, all doses  **VOGEL’S TEST:**  ↑ number of shocks, all doses | (16) |
|  | Rat, Long Evans, Both sexes  n=11/group | Vehicle vs 10 or 25 or 50 mg/kg for a single dose  Oral (gavage) | PD 27 | PD 27 & 28 | **OPEN FIELD TEST** (as part of Functional Observational Battery), same day as exposure:  ↓ locomotion starting at 25 mg/kg in males and females  ↓ rearing starting at 25 mg/kg in males and females  **OPEN FIELD TEST** (as part of Functional Observational Battery), 24 HR following exposure:  ↔ locomotion in males at all doses  ↓ locomotion starting at 50 mg/kg in females  ↔ rearing in males at all doses  ↓ rearing starting at 50 mg/kg in females | (17) |
|  | Rat, Long Evans, Both sexes  n=5/group | Vehicle vs 20 or 50 mg/kg for a single dose  Oral (gavage) | PD 27 | PD 27 | **OPEN FIELD TEST** (as part of the Functional Observational Battery), 3.5 HR following exposure:  ↓ locomotion at 20 and 50 mg/kg, both sexes  **OPEN FIELD TEST** (as part of the Functional Observational Battery), 6.5 HR following exposure:  ↓ locomotion at 20 and 50 mg/kg, males  ↓ locomotion at 50 mg/kg, females | (18) |
|  | Rat, Sprague Dawley, Both sexes  n=6-8/group | Vehicle vs 5 mg/kg/day for 34 days  Oral (gavage) | PD 27-60 | 61 | **OPEN FIELD TEST:**  ↓ locomotion | (19) |
|  | Rat, Spontaneously Hypertensive, Male  N=5/group | Vehicle vs 10 or 20 mg/kg for a single dose  Route not specified | 4 weeks | 4 weeks | **ELEVATED PLUS MAZE:**  ↔ total entries  ↔ open-arm entries | (20) |
| DFP | Rat, Sprague Dawley, Both sexes  n= 10-12/sex/group | Vehicle vs 3.4 mg/kg for a single dose  Subcutaneous  *Post treatment:*  *atropine sulfate and pralidoxime* | PD 28 | ~56 PD | **OPEN FIELD TEST:**  ↔ locomotion in both sexes | (3) |
|  | Rat, Sprague Dawley, Males  n=8-13/group | Vehicle vs 1.4 x LD50 mg/kg for a single dose  Subcutaneous  *Pre-treatment: pyridostigmine bromide;*  *Post-treatment:*  *2-PAM and atropine methylnitrate* | PD 28 | 7 and 14 months | **OPEN FIELD TEST:**  ↓ % center time at 14 months  ↔ % center time at 7 months  **ELEVATED PLUS MAZE:**  ↓ time in open arms at 7 and 14 months  **SOCIAL INTERACTION TEST:**  ↓ social contact time at 7 months and 14 months  **AGGRESSION TEST:**  ↑ aggression score at 7 and 14 months | (4) |
| Malathion | Mouse, C57BL/6, Male  n=9/group | Vehicle vs 50 mg/kg/day for 14 days  Oral (gavage) | 4-5 weeks | 6-7 weeks | **OPEN FIELD TEST:**  ↔ locomotion or time spent in zones | (6) |
| Monocrotophos | Rat, Wistar, Females  n=5/group | Vehicle vs 0.5 or 1 mg/kg/day for 27 days  Oral (gavage) | PD 22-49 | PD 50 | **OPEN FIELD TEST:**  ↓ Locomotion, all doses | (21) |
|  | Rat, Wistar, Females  n=5/group | Vehicle vs 0.5 or 1 mg/kg/day for 27 days  Oral (gavage) | PD 22-49 | PD 65 | **OPEN FIELD TEST:**  ↔ Locomotion, all doses | (21) |
| ***Pyrethroids*** | | | | | | |
| Deltamethrin | Rat, Long Evans, Both sexes  n=12/sex/dose | Vehicle vs 1, 2, or 4 mg/kg for a single dose  Oral (gavage) | PD 21 | PD 21 | **ACOUSTIC STARTLE TEST:**  ↓ response in a dose dependent manner | (22) |
|  | Rat, Wistar, Male  n=10/group | Vehicle vs 7 mg/kg/day for 15 days  Oral (gavage) | PD 22-37 | PD 38-40 | **SPONTANEOUS LOCOMOTOR ACTIVITY:**  ↑ average rate | (9) |
|  | Mouse, ICR, Female  N=3-5/group | Vehicle vs 3 or 6, or 12 mg/kg/day for 28 days  Oral | 4-8 weeks | 7 weeks | **OPEN FIELD TEST:**  ↑ locomotion in 6 mg/kg/day; ↔ in 3 and 12 mg/kg/day  ↑ center time in 6 and 12 mg/kg/day; ↔ in 3 mg/kg/day  **LIGHT/DARK ASSAY:**  ↔ time spent on dark side, all doses  **NEST BUILDING:**  ↔ nest score, all doses  **FORCED SWIM TEST:**  ↑ immobility time in 6 and 12 mg/kg/day; ↔ in 3 mg/kg/day  **CIRCADIAN ACTIVITY:** ↑ activity during day and night, 3 and 6 mg/kg/day; ↔ in 12 mg/kg/day | (23) |
|  | Rat, Wistar, Male  n=10/group | Vehicle vs 10 mg/kg for a single dose  Oral (gavage) | 8 weeks | 8 weeks (same day) | **SPONTANEOUS LOCOMOTOR ACTIVITY:**  ↔ locomotion | (24) |
| ***Neonicotinoid*** | | | | | | |
| Clothianidin | Mouse, C57Bl/6N, Both sexes  n=7-10/group | Vehicle vs 80 mg/kg for a single dose  Oral (gavage) | 6 weeks | 3 months | **OPEN FIELD TEST:**  ↔ locomotion, both sexes  ↔ time spent in center, both sexes  **LIGHT/DARK ASSAY:**  ↔ locomotion in light boxes in males, ↓ in females  ↔ latency to enter light, both sexes  ↔ transitions between light/dark, both sexes | (10) |
|  | Mouse, C57Bl/6N, Both sexes  n=6-10/group | Vehicle vs 80 mg/kg for a single dose  Oral (gavage) | 6 weeks | 7 months | **OPEN FIELD TEST:**  ↔ locomotion, both sexes  ↔ time spent in center, both sexes  **LIGHT/DARK ASSAY:**  ↔ locomotion in light boxes, both sexes  ↔ latency to enter light, both sexes  ↔ transitions between light/dark, both sexes | (10) |
| Dinotefuran | Mouse, C57BL/6NCrS1c, Male  n=6/group | Vehicle vs 100 vs 500 vs 2500 mg/kg/day for 5 weeks  Drinking water | 3-6 weeks | 8 weeks | **TAIL SUSPENSION TEST:**  ↓ immobility in 500 mg/kg/day; ↔ in 100 and 2500 mg/kg/day  **FORCED SWIM TEST:**  ↔ immobility, all doses | (25) |
| Imidacloprid | Mouse, Swiss albino, Male  n=6/group | Vehicle vs 1 or 2 mg/kg/day for 40 days  Oral (gavage) | PD 21-60 | Not specified, but before PD 60 | **OPEN FIELD TEST:**  ↔ locomotion, all doses | (11) |
| ***Organochlorine*** | | | | | | |
| Lindane | Rat, Wistar, Male  n=6/group | Vehicle vs 2.5 mg/kg/day for 21 days  Oral | 3 weeks | Not specified; presumably 6 weeks | **SPONTANEOUS LOCOMOTOR ACTIVITY:**  ↓ locomotion  ↓ rearing | (12) |

**Table S3.** Neurobiological measures (neurochemical, neuroinflammation, neurodegeneration, etc.) & Methods

| **Insecticide** | **Animal Model** | **Route & Dose** | **Exposure Age** | **Assessment Age** | **Neurobiological Outcomes** | **Reference** |
| --- | --- | --- | --- | --- | --- | --- |
| ***Organophosphates & Carbamates*** | | | | | | |
| Aldicarb | Rat, Long Evans, Both sexes  n=4/group | Vehicle vs 0.08 or 0.15 or 0.26 mg/kg for single dose  Oral (gavage) | PD 27 | PD 27 | **WHOLE BRAIN:**  Cholinergic neurotransmission  ↓ acetylcholinesterase (AChE) activity, all doses, both sexes | (13) |
| Chlorpyrifos | Rat, Sprague Dawley, Both sexes  n= 6-9/group | Vehicle vs 47 or 23.5 mg/kg for a single dose  Oral | PD 21 | PD 21 | **FRONTAL CORTEX:**  Cholinergic neurotransmission  ↓ AChE activity, all doses | (26) |
|  | Rat, Sprague Dawley, Both sexes  n= 6-9/group | Vehicle vs 47 or 23.5 mg/kg for a single dose  Oral | PD 21 | PD 22 | **FRONTAL CORTEX:**  Cholinergic neurotransmission  ↓ AChE activity, all doses | (26) |
|  | Rat, Wistar, Males  N=6/group | Vehicle vs 242 mg/kg for a single dose  Subcutaneous | PD 21 | PD 23 | **CEREBRAL CORTEX:**  Cholinergic neurotransmission  ↓ AChE activity, ↓ cholinergic acetyltransferase (ChAT), ↓ Na+, K+-ATPase  Cellular Stress & Other Intracellular Processes  ↑ lipid peroxidation levels (LPO)) | (27) |
|  | Rat, Sprague Dawley, Both sexes  n= 6-9/group | Vehicle vs 47 or 23.5 mg/kg for a single dose  Oral | PD 21 | PD 25 | **FRONTAL CORTEX:**  Cholinergic neurotransmission  ↔ AChE activity, all doses | (26) |
|  | Rat, Sprague Dawley, Sex not specified  N=3-6/group | Vehicle vs 127 mg/kg for a single dose | PD 21 | PD 25 | **FRONTO-PARIETAL CORTEX:** Cholinergic neurotransmission  ↓ AChE activity, ↔ nicotinic autoreceptor function | (28) |
|  | Rat, Sprague Dawley, Both sexes  n=6/group | Vehicle vs 38 vs 127 mg/kg for a single dose  Subcutaneous | PD 21 | PD 21-25 | **STRIATUM,** 4 HR after exposure**:**  Cholinergic neurotransmission  ↔ AChE activity, all doses  ↔ acetylcholine (ACh) synthesis, all doses  **STRIATUM,** 24 HR after exposure**:**  Cholinergic neurotransmission  ↓ AChE activity, all doses  ↔ ACh synthesis at 38 mg/kg; ↑ at 127 mg/kg  **STRIATUM,** 94 HR after exposure**:**  Cholinergic neurotransmission  ↓ AChE activity, all doses  ↔ ACh synthesis, all doses | (29) |
|  | Rat, Sprague Dawley, Both sexes  N=6-12/group | Vehicle vs 38.1 or 127 mg/kg for a single dose  Subcutaneous | PD 21 | PD 21-25 | **CEREBRAL CORTEX,** 4 HR after exposure:  Cholinergic neurotransmission  ↓ AChE activity, all doses  ↓ muscarinic receptor binding (QNB, OXO), all doses  ↓ muscarinic receptor signaling (PI hydrolysis, cAMP formation), all doses  **CEREBRAL CORTEX,** 24 HR after exposure:  Cholinergic neurotransmission  ↓ AChE activity, all doses  ↓ muscarinic receptor binding (QNB, OXO), 127 mg/kg  ↔ muscarinic receptor signaling (PI hydrolysis, cAMP formation), all doses  **CEREBRAL CORTEX,** 96 HR after exposure:  Cholinergic neurotransmission  ↓ AChE activity, all doses  ↓ muscarinic receptor binding (QNB, OXO), 127 mg/kg  ↓ muscarinic receptor signaling (PI hydrolysis, cAMP formation), 127 mg/kg | (30) |
|  | Rat, Long Evans, Both sexes  N=7-8/group | Vehicle vs 0.3 or 7 mg/kg for two doses on PD 22 and 26  Subcutaneous | PD 22 & 26 | PD 24-28 | **CORTEX:**  Cholinergic neurotransmission  ↔ AChE activity, all doses  **HIPPOCAMPUS:**  Cholinergic neurotransmission  ↔ AChE activity, all doses  **CEREBELLUM:**  Cholinergic neurotransmission  ↔ AChE activity, all doses | (1) |
|  | Rat, Sprague Dawley, Male  n=8/group | Vehicle vs 2.5 or 5 or 10 mg/kg/day for 10 days  Subcutaneous | PD 22-32 | PD 33 | **HIPPOCAMPUS:**  Cellular Stress & Other Intracellular Processes  ↑ phosphorylated GSK-3b at 10 mg/kg/day; ↔ beta-catenin; ↔ wnt2  **STRIATUM:**  Cellular Stress & Other Intracellular Processes  ↑ phosphorylated GSK-3b at 2.5, 5, and 10 mg/kg/day; ↔ beta-catenin; ↔ wnt2 | (14) |
|  | Rat, Sprague Dawley, Male | Vehicle vs 7.45 mg/kg/day for 37 days  Intraperitoneal | PD 23-60 | PD 59-60 | **RIGHT CEREBRAL HEMISPHERE:**  Neuropathology  ↑ encephalopathic changes  (e.g. necrosis, hemorrhages, gliosis)  **BRAIN** (regions not specified):  Cholinergic neurotransmission  ↓ AChE activity  Monoaminergic neurotransmission  ↓ norepinephrine  Cellular Stress & Other Intracellular Processes  ↓ total antioxidant capacity  ↑ malondialdehyde) | (15) |
|  | Rat, Sprague Dawley, Both sexes  n=5/group | Vehicle vs 20 vs 50 mg/kg for a single dose  Oral (gavage) | PD 27 | PD 27 | **WHOLE BRAIN:**  Cholinergic neurotransmission  ↓ AChE activity at 3.5 and 6.5 hours, both sexes | (18) |
|  | Rat, Sprague Dawley, Both sexes  n=6-8/group | Vehicle vs 5 mg/kg/day for 34 days  Oral (gavage) | PD 27-61 | PD 61 | **SUBSTANTIA NIGRA:**  Monoaminergic neurotransmission  ↓ Tyrosine hydroxylase (TH)  Neurodegeneration  ↑ *Stat1*, *Bax/Bcl-2* ratio, *Casp-3*, *Pkc*, *Lc3b*  **STRIATUM:**  Monoaminergic neurotransmission  ↓ 3,4-Dihydroxyphenylacetic acid (DOPAC); ↓ TH  Neurodegeneration  ↑ *Stat1*, *Bax/Bcl-2* ratio, *Casp-3*, *Pkc*, *Lc3b* | (19) |
|  | Rat, Wistar, Male  n= 6/group | Vehicle vs 80 mg/kg for a single dose  Subcutaneous | PD 28 | PD 30 | **CEREBRAL CORTEX:**  Cholinergic neurotransmission  ↓ AChE activity; ↓ choline acetyltransferase (ChAT); ↓ Na+K+ATPase; ↔ Ca2+ATPase; ↔ Mg2+ATPase  **CEREBELLUM:**  Cholinergic neurotransmission  ↓ AChE activity; ↓ ChAT; ↓ Na+K+ATPase; ↔ Ca2+ATPase; ↓ Mg2+ATPase  **MEDULLA:**  Cholinergic neurotransmission  ↓ AChE activity; ↓ ChAT; ↓ Na+K+ATPase; ↔ Ca2+ATPase; ↔ Mg2+ATPase  **SPINAL CORD:**  Cholinergic neurotransmission  ↓ AChE activity; ↓ ChAT; ↓ Na+K+ATPase; ↓ Ca2+ATPase; ↓ Mg2+ATPase) | (31) |
|  | Rat, Wistar, Male  n= 6/group | Vehicle vs 80 mg/kg for a single dose  Subcutaneous | PD 28 | PD 30 | **CEREBRAL CORTEX:**  Cellular Stress & Other Intracellular Processes  ↓ DNA content; ↓ RNA content; ↓ total protein  ↔ protein carbonyl content; ↓ total protein thiol  ↑ ALT activity; ↑ AST activity  **CEREBELLUM:**  Cellular Stress & Other Intracellular Processes  ↓ DNA content; ↓ RNA content; ↓ total protein  ↑ protein carbonyl content; ↓ total protein thiol  ↑ ALT activity; ↑ AST activity  **MEDULLA:**  Cellular Stress & Other Intracellular Processes  ↓ DNA content; ↓ RNA content; ↓ total protein  ↑ protein carbonyl content; ↓ total protein thiol  ↑ ALT activity; ↑ AST activity  **SPINAL CORD:**  Cellular Stress & Other Intracellular Processes  ↓ DNA content; ↓ RNA content; ↓ total protein  ↑ protein carbonyl content; ↓ total protein thiol  ↑ ALT activity; ↑ AST activity | (32) |
|  | Rat, Wistar, Male  n= 6/group | Vehicle vs 80 mg/kg for a single dose  Subcutaneous | PD 28 | PD 30 | **CEREBRAL CORTEX:**  Cellular Stress & Other Intracellular Processes  ↑ LPO Level; ↔ superoxide dismutase (SOD) activity; ↓ catalase (CAT) activity; ↓ glutathione peroxidase (GPx) activity; ↓ glutathione-S-transferase (GST) activity; ↓ glutathione (GSH) content  **CEREBELLUM:**  Cellular Stress & Other Intracellular Processes  ↑ LPO Level; ↔ SOD activity; ↓ CAT activity; ↓ GPx activity; ↓ GST activity; ↓ GSH content  **MEDULLA:**  Cellular Stress & Other Intracellular Processes  ↑ LPO Level; ↓ SOD activity; ↓ CAT activity; ↓ GPx activity; ↓ GST activity; ↓ GSH content  **SPINAL CORD:**  Cellular Stress & Other Intracellular Processes  ↑ LPO Level; ↓ SOD activity; ↓ CAT activity; ↓ GPx activity; ↓ GST activity; ↓ GSH content | (33) |
| Clorfenvinphos | Rat, Wistar, Both sexes  n= 10-15/sex/group | Vehicle vs 5 or 100 or 1000 ppm/day for 10 days  Oral (diet)  *Fed in two types of diet: low-protein and optimal-protein* | ~6 weeks (estimated based on reported weight) | ~7.5 weeks | **WHOLE BRAIN:**  Cholinergic neurotransmission  ↓ AChE activity at 100 and 1000 ppm, both sexes; ↔ at 5 ppm, both sexes  Cellular Stress & Other Intracellular Processes  ↓ Glucose-6-phosphate isomerase activity at 100 and 1000 ppm, both sexes, both diets; ↔ at 5 ppm in males, both diets; ↓ at 5 ppm in females, optimal-protein diet | (34) |
| DFP | Rat, Sprague Dawley, Both sexes  n = not specified | Vehicle vs 4-6.5 mg/kg for a single dose  Subcutaneous  *Pre-treatment: pyridostigmine bromide;*  *Post-treatment: atropine sulfate and 2-PAM* | PD 21 | PD 22 | **HILUS:**  Neurodegeneration  ↑ Fluoro-JadeC in animals that displayed status epilepticus  **LATERODORSAL THALAMUS:**  Neurodegeneration  ↔ Fluoro-JadeC in animals that displayed status epilepticus  **BASLOLATERAL AMYDALA:**  Neurodegeneration  ↔ Fluoro-JadeC in animals that displayed status epilepticus  **PIRIFORM CORTEX:**  Neurodegeneration  ↑ Fluoro-JadeC in animals that displayed status epilepticus  **PARIETAL CORTEX:**  Neurodegeneration  ↔ Fluoro-JadeC in animals that displayed status epilepticus | (35) |
|  | Rat, Sprague Dawley, Both sexes  n= 5-7/sex/group | Vehicle vs 3.4 mg/kg for a single dose  Subcutaneous  *Post-treatment: atropine sulfate and pralidoxime* | PD 28 | PD 29 | **CORTEX:**  Cholinergic neurotransmission  ↓ AChE activity, both sexes  **AMYGDALA:**  Neurodegeneration  ↑ Fluoro-JadeC staining, both sexes  Neuroinflammation  ↔ astrocytes & microglia, both sexes  **SOMATOSENSORY CORTEX:**  Neurodegeneration  ↑ Fluoro-JadeC staining in males, ↔ females  Neuroinflammation  ↔ astrocytes, both sexes; ↔ microglia, both sexes  **HIPPOCAMPUS:**  Neurodegeneration  ↑ Fluoro-JadeC staining, both sexes  Neuroinflammation  ↑ astrocytes in males, ↔ females; ↔ microglia, both sexes  **PIRIFORM CORTEX:**  Neurodegeneration  ↑ Fluoro-JadeC staining, both sexes  Neuroinflammation  ↔ astrocytes in both sexes; ↑ microglia in males, ↔ females  **THALAMUS:**  Neurodegeneration  ↑ Fluoro-JadeC staining, both sexes  Neuroinflammation  ↔ astrocytes, both sexes; ↑ microglia in males, ↔ females | (3) |
|  | Rat, Sprague Dawley, Both sexes  n = not specified | Vehicle vs 4-6.5 mg/kg for a single dose  Subcutaneous  *Pre-treatment: pyridostigmine bromide;*  *Post-treatment: atropine sulfate and 2-PAM* | PD 28 | PD 29 | **HILUS:**  Neurodegeneration  ↑ Fluoro-JadeC in animals that displayed status epilepticus | (35) |
|  | Rat, Sprague Dawley, Both sexes  n= 5-7/sex/group | Vehicle vs 3.4 mg/kg for a single dose  Subcutaneous  *Post-treatment: atropine sulfate and pralidoxime* | PD 28 | PD 35 | **AMYGDALA:**  Neurodegeneration  ↑ Fluoro-Jade C, both sexes  Neuroinflammation  ↑ astrocytes & ↑microglia, both sexes  **SOMATOSENSORY CORTEX:**  Neurodegeneration  ↑ Fluoro-Jade C, both sexes  Neuroinflammation  ↑ astrocytes in males, ↔ in females  ↑ microglia in males, ↔ females  **HIPPOCAMPUS:**  Neurodegeneration  ↑ Fluoro-Jade C, both sexes  Neuroinflammation  ↑ astrocytes, both sexes  ↑ microglia in males, ↔ females  **PIRIFORM CORTEX:**  Neurodegeneration  ↑ Fluoro-Jade C, both sexes  Neuroinflammation  ↑ Astrocytes in both sexes  ↑ Microglia in both sexes  **THALAMUS:**  Neurodegeneration  ↑ Fluoro-Jade C, both sexes  Neuroinflammation  ↑ astrocytes, ↑microglia, both sexes | (3) |
|  | Rat, Sprague Dawley, Both sexes  n= 5-7/sex/group | Vehicle vs 3.4 mg/kg for a single dose  Subcutaneous  *Post-treatment: atropine sulfate and pralidoxime* | PD 28 | PD 56 | **AMYGDALA:**  Neurodegeneration  ↑ Fluoro-Jade C, both sexes  Neuroinflammation  ↑ Astrocytes, both sexes  ↑ Microglia in males, ↔ females  **SOMATOSENSORY CORTEX:**  Neurodegeneration  ↑ Fluoro-Jade C, both sexes  Neuroinflammation  ↔ astrocytes, both sexes  ↑ Microglia in males, ↔ females  **HIPPOCAMPUS:**  Neurodegeneration  ↑ Fluoro-Jade C, both sexes  Neuroinflammation  ↑ Astrocytes in males, ↔ in females  ↑ Microglia in males, ↔ females  ↑ Neurogenesis in males, ↔ structural changes  **PIRIFORM CORTEX:**  Neurodegeneration  ↑ Fluoro-Jade C, both sexes  Neuroinflammation  ↑ Astrocytes in males, ↔ in females  ↑ Microglia, both sexes  **THALAMUS:**  Neurodegeneration  ↑ Fluoro-Jade C, both sexes  Neuroinflammation  ↑ Astrocytes in males, ↔ in females  ↑ Microglia, both sexes | (3) |
|  | Rat, Sprague Dawley, Males  n=6/group | Vehicle vs 1.4 x LD50 mg/kg  Subcutaneous  *Pre-treatment:*  *pyridostigmine bromide*  *Post-treatment: 2-PAM and atropine methylnitrate* | PD 28 | 14 months | **CORTEX:**  Neuroinflammation  ↑ astrocytes; ↑ microglia  Cellular composition & circuit plasticity  ↓ PV+ interneurons  **HIPPOCAMPUS:**  Neuroinflammation  ↑ astrocytes; ↑ microglia  Cellular composition & circuit plasticity  ↓ PV+ interneurons; ↓ principal neuron (NeuN+) counts; ↑ mossy fiber sprouting; ↓ neurogenesis (DCX+)  **AMYGDALA:**  Neuroinflammation  ↑ astrocytes; ↑ microglia  Cellular composition & circuit plasticity  ↓ principal neuron (NeuN+) counts | (4) |
| Diazinon | Mouse, Albino, Males  n=7/group | Vehicle vs 50 mg/kg for a single dose  Oral | ~4-5 weeks, (determined by weight) | ~7-8 weeks | **CEREBRUM** (pooled tissue together):  Cholinergic neurotransmission  ↓ AChE activity  Cellular Stress & Other Intracellular Processes  ↓ GPx; ↓ SOD; ↓ FSH; ↓ CAT  **CEREBELLUM** (pooled tissue together):  Cholinergic neurotransmission  ↓ AChE activity  Cellular Stress & Other Intracellular Processes  ↓ GPx; ↓ SOD; ↓ FSH; ↓ CAT | (36) |
| Dichlorvos | Rat, Sprague Dawley, Male  n=8-10/group | Vehicle vs 0.1 or 2 mg/kg/day for 28 days  Subcutaneous | PD 21-48 | Not specified; presumably on PD 55 or 56 following behavioral testing | **HIPPOCAMPUS:**  Neurodegeneration  ↑ Aβ_1-40_ at 0.1 mg/kg/day_,_ ↔ at 2 mg/kg/day  ↔ Aβ_1-42_ , all doses | (5) |
| Dichlofenthion | Rat, Wistar, Males  n=5/group | Vehicle vs 120 mg/kg for a single dose  Oral (stomach syringe) | 3 weeks | 3 weeks | **WHOLE BRAIN:**  Cholinergic neurotransmission  ↓ AChE activity (normalizes by 96 hours after exposure) | (37) |
|  | Rat, Wistar, Males  n=5/group | Vehicle vs 120 mg/kg for a single dose  Oral (stomach syringe) | 7 weeks | 7 weeks | **WHOLE BRAIN:**  Cholinergic neurotransmission  ↓ AChE activity (does not normalize by 168 hours after exposure) | (37) |
| Fenitrothion | Rat, Wistar, Males  n=5/group | Vehicle vs 170 mg/kg for a single dose  Oral (stomach syringe) | 7 weeks | 7 weeks | **WHOLE BRAIN:**  Cholinergic neurotransmission  ↓ AChE activity (does not normalize by 168 hours after exposure) | (37) |
| Malathion | Mouse, C57BL6, Male  N=3/group  N=2/group for transcriptomic studies | Vehicle vs 50 mg/kg/day for 14 days  Oral (gavage) | 4-5 weeks | 6-7 weeks | **HIPPOCAMPUS:**  Cellular composition & circuit plasticity  ↑ neuronal cell death (pyknotic cells), ↓ mature neurons (NeuN expression), ↓ dendritic arborization  Neuroinflammation  ↔ astrocytes  Gene regulation  ↑ transcriptomic changes (339 genes downregulated & 487 genes upregulated; top pathways: neuronal synaptic plasticity, RNA splicing, mitochondrion organization, axonogenesis)  **WHOLE BRAIN:**  Cholinergic neurotransmission  ↓ AChE activity  Cellular Stress & Other Intracellular Processes  ↓ GSH and GST | (6) |
| Monocrotophos | Rat, Wistar, Female  n=4-5/group | Vehicle vs 0.5 or 1 mg/kg/day for 27 days  Oral (gavage) | PD 22-49 | PD 50 | **FRONTAL CORTEX:**  Cholinergic neurotransmission  ↓ AChE activity, all doses  ↓ Muscarinic-cholinergic binding (H-QNB binding), all doses  **HIPPOCAMPUS:**  Cholinergic neurotransmission  ↓ AChE activity at 0.5 and 1 mg/kg, all doses  ↓ Muscarinic-cholinergic binding (H-QNB binding), all doses  **CEREBELLUM:**  Cholinergic neurotransmission  ↓ AChE activity, all doses  ↔ Muscarinic-cholinergic binding (H-QNB binding), all doses | (7) |
|  | Rat, Wistar, Female  n=4-5/group | Vehicle vs 0.5 or 1 mg/kg/day for 27 days  Oral (gavage) | PD 22-49 | PD 65 | **FRONTAL CORTEX:**  Cholinergic neurotransmission  ↓ AChE activity, all doses  ↔ Muscarinic-cholinergic binding (H-QNB binding), all doses  **HIPPOCAMPUS:**  Cholinergic neurotransmission  ↓ AChE activity at 1 mg/kg  ↔ Muscarinic-cholinergic binding (H-QNB binding) at 0.5 mg/kg/day, ↓ at 1 mg/kg/day  **CEREBELLUM:**  Cholinergic neurotransmission  ↔ AChE activity, both doses  ↔ Muscarinic-cholinergic binding (H-QNB binding), both doses | (7) |
|  | Rat, Wistar, Females | Vehicle vs 0.5 or 1 mg/kg/day for 27 days  Oral (gavage) | PD 22-49 | PD 50 | **FRONTAL CORTEX:**  Monoaminergic neurotransmission  ↑ 5-HT2A receptors at 1 mg/kg/day  Cellular Stress & Other Intracellular Processes  ↑ LPO, both doses; ↑ Protein carbonyls at 1 mg/kg/day; ↓ GSH, both doses; ↓ GSH peroxidase, both doses; ↓ SOD, 1 mg/kg/day; ↓ CAT, both doses  **HIPPOCAMPUS:**  Cellular Stress & Other Intracellular Processes  ↔ LPO both doses; ↑ protein carbonyls, both doses; ↓ GSH, both doses; ↓ GSH peroxidase, both doses; ↓ SOD, both doses; ↓ CAT, 1 mg/kg/day  **STRIATUM:**  Monoaminergic neurotransmission  ↓ DA-D2 receptors, both doses  Cellular Stress & Other Intracellular Processes  ↑ Oxidative Stress at 0.5 and 1 mg/kg/day; ↑ LPO, both doses; ↑ protein carbonyls, both doses; ↓ GSH, both doses; ↓ GSH peroxidase, both doses; ↓ SOD, both doses; ↓ CAT, both doses  **CEREBELLUM:**  Cellular Stress & Other Intracellular Processes  ↑ LPO at 1 mg/kg/day; ↑ protein carbonyls at 1 mg/kg/day; ↓ GSH, both doses; ↓ GSH peroxidase, both doses; ↓ SOD, both doses; ↓ CAT, both doses | (21) |
|  | Rat, Wistar, Females | Vehicle vs 0.5 or 1 mg/kg/day for 27 days  Oral (gavage) | PD 22-49 | PD 65 | **FRONTAL CORTEX:**  Monoaminergic neurotransmission  ↑ 5-HT2A receptors at 1 mg/kg/day  Cellular Stress & Other Intracellular Processes  ↔ LPO, both doses; ↑ protein carbonyls, both doses; ↓ GSH, 1 mg/kg/day; ↓ GSH peroxidase, both doses; ↓ SOD, 1 mg/kg/day; ↓ CAT, 1 mg/kg/day  **STRIATUM:**  Monoaminergic neurotransmission  ↓ DA-D2 receptors at 0.5 and 1 mg/kg/day  Cellular Stress & Other Intracellular Processes  ↑ LPO at 1 mg/kg/day; ↔ protein carbonyls, all doses; ↓ GSH, all doses; ↓ GSH peroxidase, all doses; ↓ SOD, all doses; ↓ CAT, all doses  **HIPPOCAMPUS:**  Cellular Stress & Other Intracellular Processes  ↑ LPO at 1 mg/kg/day; ↑ protein carbonyls, all doses; ↔ GSH, both doses; ↓ GSH peroxidase, all doses; ↓ SOD, 1 mg/kg/day; ↔ CAT, both doses  **CEREBELLUM:**  Cellular Stress & Other Intracellular Processes  ↔ LPO, all doses; ↑ protein carbonyls at 1mg/kg/day; ↓ GSH, all doses; ↓ GSH peroxidase, all doses; ↔ SOD, all doses; ↔ CAT, both doses | (21) |
| Parathion | Rat, Sprague Dawley, Both sexes  n=6/group | Vehicle vs 1.44 vs 4.8 mg/kg for a single dose  Subcutaneous | PD 21 | PD 21-25 | **STRIATUM,** 4 HR following exposure**:**  Cholinergic neurotransmission  ↓ AChE activity at 1.44 mg/kg, ↓ at 4.8 mg/kg  ↔ AChE synthesis at 1.44 mg/kg, ↑ at 4.8 mg/kg  **STRIATUM,** 24 HR following exposure**:**  Cholinergic neurotransmission  ↓ AChE activity, both doses  ↔ AChE synthesis at 1.44 mg/kg, ↑ at 4.8 mg/kg  **STRIATUM,** 94 HR following exposure**:**  Cholinergic neurotransmission  ↔ AChE activity at 1.44 mg/kg, ↓ at 4.8 mg/kg  ↔ AChE synthesis, all doses | (29) |
|  | Rat, Sprague Dawley, Male  n=6/group | Vehicle vs 6 mg/kg for a single dose  Subcutaneous | PD 24 | PD 26, 28, 31 | **FRONTAL CORTEX:**  Cholinergic neurotransmission  ↓ AChE activity | (38) |
|  | Rat, Sprague Dawley, Male  n=6/group | Vehicle vs 6 mg/kg for a single dose  Subcutaneous | PD 24 | PD 28 | **FRONTAL CORTEX:**  Cholinergic neurotransmission  ↓ AChE activity | (38) |
|  | Rat, Sprague Dawley, Male  n=6/group | Vehicle vs 6 mg/kg for a single dose  Subcutaneous | PD 24 | PD 31 | **FRONTAL CORTEX:**  Cholinergic neurotransmission  ↓ AChE activity | (38) |
|  | Rat, Long Evans, Males  n=6/group | Vehicle vs 100 ug/kg for a single dose  Intramuscular | PD 25-30 | PD 25-30 | **CORTEX:**  Neuropathology  ↑ leaky capillaries (horseradish peroxidase)  ↑ extravasation of albumin (into parenchyma, compared to location inside lumen in controls)  Cholinergic neurotransmission  ↓ AChE activity  **STRIATUM:**  Neuropathology  ↑ leaky capillaries (horseradish peroxidase)  ↑ extravasation of albumin (into parenchyma, compared to location inside lumen in controls)  **HIPPOCAMPUS:**  Neuropathology  ↑ leaky capillaries (horseradish peroxidase)  **TEMPORAL CORTEX:**  Cholinergic neurotransmission  ↓ AChE activity | (39) |
| Trichlorfon | Rat, Wistar, Males  n=5/group | Vehicle vs 310 mg/kg for a single dose  Oral (stomach syringe) | 7 weeks | 7 weeks | **WHOLE BRAIN:**  Cholinergic neurotransmission  ↓ AChE activity (normalizes by 48 hours after exposure) | (37) |
| ***Pyrethroids*** | | | | | | |
| Cypermethrin | Rat, Wistar, Males  n=5/group | Vehicle vs 10 or 25 mg/kg/day for 2 weeks  Oral (gavage) | PD 24 | PD 38 | **CORTEX:**  Neurodegeneration  ↑ Amyloid/Tau pathology (Aβ1-42 levels, p-tau) at 25 mg/kg/day; ↔ at 10 mg/kg/day  **HIPPOCAMPUS:**  Neurodegeneration  ↑ Amyloid/Tau pathology (Aβ1-42 levels, p-tau) at 25 mg/kg/day; ↔ at 10 mg/kg/day | (8) |
|  | Rat, Wistar, Males  n=5/group | Vehicle vs 10 or 25 mg/kg/day for 3 weeks  Oral (gavage) | PD 24 | PD 45 | **CORTEX:**  Neurodegeneration  ↑ Amyloid/Tau pathology (Aβ, p-tau, tau, APP, PS1)  ↑ APP cleavage and tau phosphorylation pathways (APP, PS1, PS2, GSK3β)  ↓ HB-EGF Signaling  Neuroinflammation  ↑ IL-1β, NF-κB, GFAP, IL-1R1  **HIPPOCAMPUS:**  Neurodegeneration  ↑ Amyloid/Tau pathology (Aβ, p-tau, tau, APP, PS1), both groups  ↑ APP cleavage and tau phosphorylation pathways (APP, PS1, GSK3β)  ↓ HB-EGF Signaling  Neuroinflammation  ↑ IL-1β, NF-κB, GFAP, IL-1R1 | (8) |
| Deltamethrin | Rat, Wistar, Male  n=10/group | Vehicle vs 10 mg/kg for a single dose  Oral | 8 weeks | 8 weeks (same day) | **WHOLE BRAIN:**  Cellular Stress & Other Intracellular Processes  ↔ P450 monooxygenases (EROD and PROD) | (24) |
|  | Rat, Wistar, Male  n=5/group (different set of mice for each measured outcome) | Vehicle vs 7 mg/kg/day for 15 days  Oral (gavage) | PD 22-37 | PD 38 | **WHOLE BRAIN:**  Monoaminergic neurotransmission  ↑ MAO  Cholinergic Neurotransmission  ↔ Na^+^K^+^-ATPase  ↑ AChE activity  **FRONTAL CORTEX:**  Cellular Stress & Other Intracellular Processes  ↔ putrescine, ↑ spermidine, ↑ spermine)  **CORPUS STRIATUM:**  Cellular Stress & Other Intracellular Processes  ↑ putrescine, ↑ spermidine, ↑ spermine  Monoaminergic neurotransmission  ↑ Dopamine receptor binding  ↓ Muscarinic receptor binding  **HIPPOCAMPUS:**  Cellular Stress & Other Intracellular Processes  ↑ putrescine, ↓ spermidine, ↓ spermine  **CEREBELLUM:**  Cellular Stress & Other Intracellular Processes  ↓ putrescine, ↓ spermidine, ↓ spermine  **HYPOTHALAMUS:**  Cellular Stress & Other Intracellular Processes  ↑ putrescine, ↑ Spermidine, ↑ Spermine  **PONS MEDULLA:**  Cellular Stress & Other Intracellular Processes  ↔ putrescine, ↓ spermidine, ↓ spermine | (9) |
|  | Mouse, *Arc-Luc* Tg, homogyzgous, Sex not specified  n= 3-4/group | Vehicle vs 3.5 mg/kg for a single dose  Intraperitoneal | 4 weeks | 4 weeks (same day) | **Bioluminescence signal in this strain of mice is interpreted as neuronal activity by this study*  **FOREBRAIN:**  Cellular Stress & Other Intracellular Processes  ↔ bioluminescence at 3 or hours following exposure | (40) |
|  | Mouse, *Arc-Luc* Tg. Heterozygous, Sex not specified  n=5-6 | Vehicle vs 8 mg/kg every other day for 4 weeks  Oral (gavage) | PD 28-56 | PD 28, 35, 42, 49, & 56 | **Bioluminescence signal in this strain of mice is interpreted as neuronal activity by this study*  **FOREBRAIN:**  Cellular Stress & Other Intracellular Processes  ↔ bioluminescence at PD28, 35, 42, 49, and 56 | (40) |
|  | Mouse, ICR, Female  N=5/group | Vehicle vs 3 or 6, or 12 mg/kg/day for 28 days  Oral (gavage) | 4-8 weeks | 7 weeks | **HYPOTHALAMUS:**  Neuroendocrine systems  ↓ thyroid hormone homeostasis gene (*Mct8*), all groups; ↓ *Dio2* in 6 and 12 mg/kg/day  **PITUITARY:**  Neuroendocrine systems  ↓ thyroid hormone homeostasis gene (*dio2*), all groups; ↔ *Mct8* and *Trhr* in all groups  **STRIATUM:**  Monoaminergic neurotransmission  ↑ DAT protein, all doses  ↑ TH protein, all doses | (23) |
| ***Neonicotinoid*** | | | | | | |
| Dinotefuran | Mouse, C57BL/6NCrS1c, Male  n=6/group | Vehicle vs 100 or 500 or 2500 mg/kg/day for 5 weeks  Drinking water | 3-8 weeks | 8 weeks | **DORSAL RAPHE NUCLEUS:**  Monoaminergic neurotransmission  ↔ 5-HT, all exposures | (25) |
| Imidacloprid | Mouse, Swiss albino, Male  n=6/group | Vehicle vs 1 or 2 mg/kg/day for 40 days  Oral (gavage) | PD 21-60 | PD 60 | **WHOLE BRAIN:**  Cholinergic neurotransmission  ↔ AChE in 1 and 2 mg/kg/day  Cellular Stress & Other Intracellular Processes  ↓ CAT in 1 mg/kg/day, ↔ at 2 mg/kg/day  ↔ SOD  ↑ LPO in 2 mg/kg/day, ↔ at 1 mg/kg/day | (11) |
|  | Mouse, *Arc-Luc* Tg, homogyzgous  n= 3-4/group | Vehicle vs 40 mg/kg for a single dose  Intraperitoneal | 4 weeks | 4 weeks (same day) | **Bioluminescence signal in this strain of mice is interpreted as neuronal activity by this study*  **FOREBRAIN:**  Cellular Stress & Other Intracellular Processes  ↔ bioluminescence at 3 or hours following exposure | (40) |
| Clothianidin | Mouse, C57Bl/6N, Both sexes  n=6-10/group | Vehicle vs 80 mg/kg for a single dose  Oral (gavage) | 6 weeks | 7 months | **HIPPOCAMPUS:**  Neuroinflammation  ↔ astrocytes (GFAP), both sexes  Cellular composition & circuit plasticity  ↔ neural stem cells (SOX2), both sexes  ↔ immature neurons (DCX), both sexes | (10) |
| ***Organochlorine*** | | | | | | |
| Chlordecone | Mouse, ICR, Male  n=5/group | Vehicle vs 25 mg/kg for a single dose  Oral | 6-8 weeks | 6-8 weeks  (24 hours following exposure) | **WHOLE BRAIN:**  Cellular Stress & Other Intracellular Processes  ↑ total calcium, ↑ protein-bound calcium  **WHOLE BRAIN** (subcellular fraction):  Cellular Stress & Other Intracellular Processes  ↑ calcium in whole homogenate, ↑ calcium in nuclear, ↑ calcium in crude mitochondrial, ↔ calcium in microsomal, ↔ calcium in synaptosome, ↑ calcium in myelin, ↔ calcium in supernatant | (41) |
|  | Mouse, ICR, Male  n=3/group | Vehicle vs 25 mg/kg/day for 8 days  Oral | 6-8 weeks until ~7-9 weeks | ~7-9 weeks (24 hours following exposure) | **WHOLE BRAIN:**  Cellular Stress & Other Intracellular Processes  ↓ total calcium, ↓ protein-bound calcium  **WHOLE BRAIN** (subcellular fraction):  Cellular Stress & Other Intracellular Processes  ↑ calcium in whole homogenate, ↑ calcium in nuclear, ↑ calcium in crude mitochondrial, ↔ calcium in microsomal, ↔ calcium in synaptosome, ↑ calcium in myelin, ↔ calcium in supernatant | (41) |
| DDT | Rat, Wistar, Male  n=6/group | Vehicle vs 0.006, 0.06, 0.6, 6, 60 mg/kg/day for 4 weeks  Oral (gavage) | 3 weeks | 7 weeks | **CEREBRUM:**  Cellular Stress & Other Intracellular Processes  ↓ LPO in 0.06 and 0.6 mg/kg/day | (42) |
|  | Rat, Wistar, Male  n=6/group | Vehicle vs 0.06 mg/kg/day for 4 weeks  Oral (gavage) | 3 weeks | 7 weeks | **HIPPOCAMPUS:**  Gene Regulation  ↔ gene expression alterations in microarray  **HYPOTHALAMUS:**  Gene Regulation  ↑ 40 genes altered in microarray  ↑ 6 CpG island hypomethylation in MeD-PCR | (42) |
|  | Rat, Wistar, Male  n=6/group | Vehicle vs 0.06 vs 60 mg/kg/day for 4 weeks  Oral (gavage) | 3 weeks | 7 weeks | **HYPOTHALAMUS:**  Gene Regulation  ↔ global DNA methylation | (42) |
| Hepatochlor | Rat, Sprague Dawley, Both sexes  n= 4/sex/group | Vehicle vs 3 mg/kg/day for 21 days  Oral (gavage) | PD 21-42 | PD 43 | **STRIATUM:**  Monoaminergic neurotransmission  ↑ Dopamine transporter binding in males; ↔ in females | (43) |
|  | Rat, Sprague Dawley, Both sexes  n= 4/sex/group | Vehicle vs 3 mg/kg/day for 21 days  Oral (gavage) | PD 21-42 | PD 128 | **STRIATUM:**  Monoaminergic neurotransmission  ↑ Dopamine transporter binding, both sexes | (43) |
| Lindane | Rat, Wistar, Male  n=6/group | Vehicle vs 2.5 mg/kg/day for 21 days  Oral | 3 weeks | Not specified; presumably 6 weeks | **HIPPOCAMPUS:**  Cellular Stress & Other Intracellular Processes  ↑ reactive oxygen species (ROS), ↑ LPO, ↓ glutathione reductase (GR) content, ↓ SOD activity, ↓ CAT activity  ↓ ubiquitin proteasome pathway associated proteins (UBC, UCHL1, UBCH5, UBE1A, UBE2L3, UBE2N)  ↑ Cytochrome C in cytosol; ↓ in mitochondria  Cholinergic neurotransmission  ↑ AChE, ChAT, CHRM2  Neurodegeneration  ↑ Parkinson’s associated proteins (Parkin, PINK1, SNCA)  ↑ Alzheimer’s associated proteins (APP, Aβ42, Tau, pTau, Bace1)  ↑ autophagy associated proteins (Beclin-1, Atg 5, Atg 12, LC3 a, LC3 b)  ↑ apoptosis associated proteins (Bax, Bad, ↓ BC12, P53, Caspase 3, Caspase 9)  *Altered protein levels from proteomics (selected list; see original study for full list):*   - *Cytoskeleton/Structural*: ↓ β-tubulin (2A, 2B, 3A, 4B), ↓ α-tubulin 1A, ↓ stathmin, ↑ calponin-3, ↑ GFAP - *Synaptic/Signaling*: ↑ calmodulin, ↑ α-synuclein, ↑ Septin-5, ↑ 14-3-3ε, ↑ 14-3-3ζ/δ, ↓ endophilin A1, ↓ dynactin-2, ↓ profilin-2 - *Proteostasis (UPS):* ↑ proteasome subunit β6, ↑ proteasome subunit α5, ↑ proteasome subunit B4, ↓ ubiquitin conjugating enzyme, ↓ UCHL-1, ↓ UBA5 - *Antioxidant/Stress*: ↓ Cu-Zn SOD, ↓ peroxiredoxin-2, ↓ peroxiredoxin-6, ↓ lactoglutathion lyase, ↓ DJ-1/Park7 - *Metabolic/Mitochondrial*: ↓ ATP synthase β, ↓ creatine kinase, ↓ pyruvate dehydrogenase E1β, ↓ isocitrate dehydrogenase a, ↓ malate dehydrogenase-1, ↓ LDH-B, ↓ 3-mercaptopyruvate sulphotransferase - *Heat Shock/Chaperones*: ↓ HSP-90α, ↓ HSC-71, ↓ HSP-60, ↑ stress-70 - *Other*: ↑ APOE precursor, ↑ cathepsin B, ↑ tau tubulin kinase, ↑ α-enolase, ↑ γ-enolase, ↓ PDIA3, ↓ guanine deaminase   **SUBSTANTIA NIGRA:**  Cellular Stress & Other Intracellular Processes  ↑ ROS, ↑ LPO, ↓ GR content, ↓ SOD activity, ↓ CAT activity  ↓ ubiquitin proteasome pathway associated proteins (UBC, UCHL1, UBCH5, UBE1A, UBE2L3, UBE2N)  ↑ Cytochrome C in cytosol; ↓ in mitochondria  Monoaminergic Neurotransmission  ↑ neurotransmission proteins (TH, DAD2)  Neurodegeneration  ↑ Parkinson’s associated proteins (Parkin, PINK1, SNCA)  ↑ Alzheimer’s associated proteins (APP, Aβ42, Tau, pTau, Bace1)  ↑ autophagy associated proteins (Beclin-1, Atg 5, Atg 12, LC3 a, LC3 b)  ↑ apoptosis associated proteins (Bax, Bad, ↓ BC12, P53, Caspase 3, Caspase 9)  *Altered protein levels from proteomics (selected list; see original study for full list):*   - *Cytoskeleton/Structural*: ↓ β-tubulin (2A, 2B, 3A, 4B), ↓ α-tubulin 1A, ↓ stathmin, ↑ calponin 3, ↑ GFAP - *Synaptic/Signaling*: ↑ calmodulin, ↑ α-synuclein, ↑ Septin-5, ↑ 14-3-3ε, ↑ 14-3-3ζ/δ, ↓ endophilin A1, ↓ dynactin subunit-2, ↓ profilin-2, ↓ guanine nucleotide-binding protein β-1 - *Proteostasis/Ubiquitin–Proteasome System*: ↑ proteasome subunit β6, ↑ proteasome subunit α5, ↑ proteasome subunit B4, ↓ ubiquitin conjugating enzyme, ↓ UCHL-1, ↓ ubiquitin-like modifier–activating enzyme 5 - *Oxidative Stress / Antioxidants*: ↓ Cu-Zn SOD, ↓ peroxiredoxin-2, ↓ peroxiredoxin-6, ↓ lactoglutathion lyase, ↓ DJ-1/Park7, ↓ guanine deaminase - *Metabolic / Mitochondrial Enzymes*: ↓ ATP synthase β-subunit, ↓ brain creatine kinase, ↓ pyruvate dehydrogenase E1β, ↓ isocitrate dehydrogenase NAD subunit a, ↓ malate dehydrogenase-1, ↓ 3-mercaptopyruvate sulphotransferase, ↓ inosine triphosphate pyrophosphatase - *Heat Shock / Chaperones*: ↓ HSP-90α, ↓ HSC-71, ↓ HSP-60, ↑ stress-70, ↓ T-complex protein 1β, ↓ PDIA3 - *Other / Miscellaneous*: ↑ APOE precursor, ↑ cathepsin B, ↑ tau tubulin kinase, ↑ α-enolase, ↑ γ-enolase, ↓ glial maturation factor-B, ↓ dihydropyrimidinase-related protein-2, ↓ V-type proton ATPase catalytic subunit A (VATA), ↓ phosphatidyl ethanolamine binding protein 1 & 2 | (12) |
|  | Rat Wistar, Both sexes  n=4/group/sex | Vehicle vs 20 mg/kg for a single dose  Oral (gavage) | PD 22 | PD 22  (1 hr after) | **FRONTAL CORTEX:**  Monoaminergic neurotransmission  ↔ Noradrenaline, ↔ Serotonin (5-HT), ↔ 5HIAA, ↓ 5-HIAA/5-HT concentration ratio  **CORPUS STRIATUM:**  Monoaminergic neurotransmission  ↔ Noradrenaline, ↔ Dopamine, ↔ DOPAC, ↔ Homovanillic acid (HVA), ↔ DOPAC/DA concentration ratio, ↔ Serotonin (5-HT), ↔ 5-HIAA  **HIPPOCAMPUS:**  Monoaminergic neurotransmission  ↔ Noradrenaline, ↔ 5-HT, ↔ 5HIAA, ↔ 5-HIAA/5-HT concentration ratio  **HYPOTHALAMUS:**  Monoaminergic neurotransmission  ↔ Noradrenaline, ↔ Dopamine, ↔ DOPAC, ↔ Serotonin (5-HT), ↔ 5-HIAA  **THALAMUS:**  Monoaminergic neurotransmission  ↔ Noradrenaline, ↔ Dopamine, ↔ Serotonin (5-HT), ↔ 5-HIAA  **MESENCEPHALON:**  Monoaminergic neurotransmission  ↓ Noradrenaline, ↔ Dopamine, ↔ 3,4-dyhydroxyphenyleactic acid (DOPAC), ↔ Homovanillic acid (HVA), ↔ DOPAC/DA concentration ratio**,** ↔ Serotonin (5-HT), ↔ 5-HIAA  **PONS/MEDULLA:**  Monoaminergic neurotransmission  ↔ Noradrenaline, ↔ Serotonin (5-HT), ↔ 5-HIAA, ↔ 5-HIAA/5-HT concentration ratio  **COLLICULI:**  Monoaminergic neurotransmission  ↔ Noradrenaline, ↔ Serotonin (5-HT), ↔ 5-HIAA, ↔ 5-HIAA/5-HT concentration ratio | (44) |
|  | Rat, Wistar, Both sexes  n=4-5/group | Vehicle vs 10 mg/kg/day for 7 days  Oral (gavage) | PD 22-28 | PD 29 | *Summary of main findings, only. This study reports for each subregion within the broader regions. This study measured 2-DG uptake as a marker for local functional neural alterations.*  **CORTEX:**  Metabolism & Cellular Stress (Energy Metabolism)  ↔ 2-DG uptake  **BASAL GANGLIA:**  Metabolism & Cellular Stress (Energy Metabolism)  ↔ 2-DG uptake  **SEPTUM:**  Metabolism & Cellular Stress (Energy Metabolism)  ↔ 2-DG uptake  **HIPPOCAMPUS:**  Metabolism & Cellular Stress (Energy Metabolism)  ↓ 2-DG uptake  **HYPOTHALAMUS:**  Metabolism & Cellular Stress (Energy Metabolism)  ↔ 2-DG uptake  **THALAMUS:**  Metabolism & Cellular Stress (Energy Metabolism)  ↔ 2-DG uptake  **BRAINSTEM:**  Metabolism & Cellular Stress (Energy Metabolism)  ↓ 2-DG uptake  **CEREBELLUM:**  Metabolism & Cellular Stress (Energy Metabolism)  ↓ 2-DG uptake | (45) |
|  | Rat, Wistar, Both sexes  n=4/group/sex | Vehicle vs 20 mg/kg for a single dose  Oral (gavage) | PD 29 | PD 29 | **FRONTAL CORTEX:**  Monoaminergic neurotransmission  ↔ Noradrenaline, ↔ Serotonin (5-HT), ↔ 5-HIAA, ↑ 5-HIAA/5-HT concentration ratio  **CORPUS STRIATUM:**  Monoaminergic neurotransmission  ↔ Noradrenaline, ↔ Serotonin (5-HT), ↔ 5-HIAA, ↔ Dopamine, ↔ DOPAC, ↔ Homovanillic acid (HVA), ↔ DOPAC/DA concentration ratio  **HIPPOCAMPUS:**  Monoaminergic neurotransmission  ↔ Noradrenaline, ↔ Serotonin (5-HT), ↔ 5-HIAA, ↔ 5-HIAA/5-HT concentration ratio  **HYPOTHALAMUS:**  Monoaminergic neurotransmission  ↔ Noradrenaline, ↔ Serotonin (5-HT), ↔ 5-HIAA, ↔ Dopamine, ↔ DOPAC  **THALAMUS:**  Monoaminergic neurotransmission  ↔ Noradrenaline, ↔ Serotonin (5-HT), ↔ 5-HIAA. ↔ Dopamine, ↔ DOPAC, DOPAC/DA concentration ratio  **MESENCEPHALON:**  Monoaminergic neurotransmission  ↔ Noradrenaline, ↔ Serotonin (5-HT), ↔ 5-HIAA, ↔ Dopamine (DA), ↔ DOPAC, ↑ DOPAC/DA concentration ratio  **PONS/MEDULLA:**  Monoaminergic neurotransmission  ↔ Noradrenaline, ↔ Serotonin (5-HT), ↔ 5-HIAA, ↑ 5-HIAA/5-HT concentration ratio  **COLLICULI:**  Monoaminergic neurotransmission  ↓ Noradrenaline, ↔ Serotonin (5-HT), ↔ 5-HIAA, ↑ 5-HIAA/5-HT concentration ratio | (44) |
|  | Rat, Wistar, Both sexes  n=4-10/group | Vehicle vs 10 or 20 mg/kg for a single dose  Oral (gavage) | PD 29 | PD 29 | *Summary of main findings, only. This study reports for each subregion within the broader regions. This study measured 2-DG uptake as a marker for local functional neural alterations.*  **CORTEX:**  Cellular Stress & Other Intracellular Processes  ↔ 2-DG uptake  **BASAL GANGLIA:**  Cellular Stress & Other Intracellular Processes  ↑ 2-DG uptake at 20 mg/kg  **SEPTUM:**  Cellular Stress & Other Intracellular Processes  ↔ 2-DG uptake  **HIPPOCAMPUS:**  Cellular Stress & Other Intracellular Processes  ↔ 2-DG uptake  **HYPOTHALAMUS:**  Cellular Stress & Other Intracellular Processes  ↔ 2-DG uptake  **THALAMUS:**  Cellular Stress & Other Intracellular Processes  ↑↑ 2-DG uptake at 20 mg/kg  **BRAINSTEM:**  Cellular Stress & Other Intracellular Processes  ↓ 2-DG uptake at 20 mg/kg  **CEREBELLUM:**  Cellular Stress & Other Intracellular Processes  ↔ 2-DG uptake | (45) |

**Table S4.** General neurotoxic outcomes (seizures, tremors, neuromuscular function alterations) & Methods

| **Insecticide** | **Animal Model** | **Route & Dose** | **Exposure Age** | **Assessment Age** | **Effects on Neurotoxic Outcomes** | **Reference** |
| --- | --- | --- | --- | --- | --- | --- |
| ***Organophosphates & Carbamates*** | | | | | | |
| Aldicarb | Rat, Long Evans, Both sexes  n=10/group | Vehicle vs 0.08 or 0.15 or 0.26 mg/kg for a single dose  Oral (gavage) | PD 27 | PD 27 & 28 | **FUNCTIONAL OBSERVATIONAL BATTERY**, 1 HR following exposure:  ↓ arousal in 0.15 and 0.26 mg/kg, both sexes  ↓ handling reactivity in 0.26 mg/kg, both sexes  ↓ righting reflex in 0.15 and 0.26 mg/kg, both sexes  ↓ tail-pinch response, all doses, in males, ↓ at 0.26 mg/kg, females  **FUNCTIONAL OBSERVATIONAL BATTERY**, 24 HR following exposure:  ↔ all measures, all doses, both sexes | (13) |
| Chlorpyrifos | Rat, Long Evans, Both sexes  N=7-8/group | Vehicle vs 0.3 or 7 mg/kg for two doses  Subcutaneous | PD 22 and 26 | PD 24 & 28 | ↔ tremors, altered gait, and excessive excretions | (1) |
|  | Rat, Long Evans, Both sexes  n=11/group | Vehicle vs 10 or 25 or 50 mg/kg for a single dose  Oral (gavage) | PD 27 | PD 27 & 28 | **FUNCTIONAL OBSERVATIONAL BATTERY**, same day as exposures:  ↓ gait/ataxia starting at 25 mg/kg, both sexes  ↓ righting reflex starting at 50 mg/kg, males and at 25 mg/kg, females  ↓ arousal starting at 50 mg/kg, males and at 25 mg/kg, females  ↓ tail-pinch response starting at 50 mg/kg, males and at 25 mg/kg, females  ↑ tremors starting at 25 mg/kg, both sexes  ↑ lacrimation starting at 50 mg/kg, both sexes  **FUNCTIONAL OBSERVATIONAL BATTERY**, 24 HR following exposures:  ↓ gait/ataxia starting at 50 mg/kg, both sexes  ↓ righting reflex starting at 50 mg/kg, females  ↔ arousal at all doses, both sexes  ↔ tail-pinch response at all doses, both sexes  ↔ tremors for all doses in males, ↑ starting at 50 mg/kg, females  ↔ lacrimation for all doses, both sexes | (17) |
|  | Rat, Long Evans, Both sexes  n=5/group | Vehicle vs 20 or 50 mg/kg for a single dose  Oral (gavage) | PD 27 | PD 27 | **FUNCTIONAL OBSERVATIONAL BATTERY**, 3.5 HR following exposures:  ↓ gait/ataxia at 50 mg/kg, both sexes  ↑ tremors starting at 50 mg/kg, both sexes  ↑ lacrimation starting at 50 mg/kg, both sexes  ↓ tail-pinch response at 50 mg/kg, males  ↓ tail-pinch response at 20 and 50 mg/kg, females  **FUNCTIONAL OBSERVATIONAL BATTERY**, 6.5 HR following exposures:  ↔ gait/ataxia, both doses, males  ↓ gait/ataxia at 50 mg/kg, females  ↔ tremors, both doses, both sexes  ↔ lacrimation, both doses, both sexes  ↓ tail-pinch response at 50 mg/kg, both sexes | (18) |
| DFP | Rat, Sprague Dawley, Both sexes  n=82 total | Vehicle vs 4.5-6 mg/kg for a single dose  Subcutaneous  *Pre-treatment: pyridostigmine bromide;*  *Post-treatment: atropine sulfate and 2-PAM* | PD 21 | PD 21-22 | ↑ Seizures (confirmed status epilepticus, spontaneous bursts of seizure activity, or interictal spiking by EEG) | (35) |
|  | Rat, Sprague Dawley, Both sexes  n= 15/sex | 3.4 mg/kg for a single dose  Subcutaneous  *Post-treatment: atropine sulfate and pralidoxime* | PD 28 | PD 28 | ↑ Seizures and ↑ seizure duration, more severe in males (supported by EEG) | (3) |
|  | Rat, Sprague Dawley, Both sexes  n=46 total | Vehicle vs 4-6.5 mg/kg for a single dose  Subcutaneous  *Pre-treatment: pyridostigmine bromide;*  *Post-treatment: atropine sulfate and 2-PAM* | PD 28 | PD 28-29 | ↑ Seizures (confirmed status epilepticus, spontaneous bursts of seizure activity, or interictal spiking by EEG) | (35) |
| Fenthion | Mouse, Albino Swiss webster, Males  N=3 | 600 mg/kg for a single dose  Intraperitoneal | 1 month | 1 month | ↑ Clinical neurotoxic signs (level 4 = “unable to walk, legs cannot support the body, difficulty breathing”) | (46) |
| Monocroptophos | Rat, Wistar, Female  n=4-5/group | Vehicle vs 0.5 or 1 mg/kg/day for 27 days  Oral (gavage) | PD 22-49 | PD 50 | ↓ Grip strength at 0.5 mg/kg/day and 1 mg/kg/day | (7) |
|  | Rat, Wistar, Female  n=4-5/group | Vehicle vs 0.5 or 1 mg/kg/day for 27 days  Oral (gavage) | PD 22-49 | PD 65 | ↓ Grip strength at 0.5 mg/kg/day and 1 mg/kg/day | (7) |
|  | Rat, Wistar, Females  n=5/group | Vehicle vs 0.5 or 1 mg/kg/day for 27 days  Oral (gavage) | PD 22-49 | PD 50 | **ROTAROD:**  ↓ time to fall, all doses | (21) |
|  | Rat, Wistar, Females  n=5/group | Vehicle vs 0.5 or 1 mg/kg/day for 27 days  Oral (gavage) | PD 22-49 | PD 65 | **ROTAROD:**  ↓ time to fall at 1 mg/kg/day; ↔ at 0.5 mg/kg/day | (21) |
| Paraoxon | Rat, Long Evans, Male  n=6/group | Vehicle vs 100 ug/kg paraoxon for a single dose  Intramuscular | PD 25-30 | PD 25-30 | ↔ SLUD signs (salivation, lacrimation, urination, and defecation)  ↔ involuntary movements | (39) |
| ***Pyrethroids*** | | | | | | |
| Deltamethrin | Rat, Sprague Dawley, Male  n=4/group | Vehicle vs 10 mg/kg for a single dose  Oral (gavage) | PD 21 | PD 21 | ↑ salivations from 1-6 hours  ↑ tremors from 1-6 hours | (47) |
|  | Rat, Sprague Dawley, Male  n not specified | Vehicle vs 2 or 10 mg/kg for a single dose  Oral (gavage) | PD 21 | PD 21 | ↑ salivations, tremors, choreoathetosis at all doses | (48) |
|  | Rat, Sprague Dawley, Male  n=4/group | Vehicle vs 10 mg/kg for a single dose  Oral (gavage) | PD 40 | PD 40 | ↑ salivations from 1-3 hours  ↔ tremors | (47) |
|  | Rat, Sprague Dawley, Male  n not specified | Vehicle vs 2 or 10 mg/kg for a single dose  Oral (gavage) | PD 40 | PD 40 | ↔ salivations, tremors at 2 mg/kg  ↑ salivation (mild-to-moderate) at 10 mg/kg | (48) |
|  | Rat, Wistar, Male  n=10/group | Vehicle vs 10 mg/kg for a single dose  Oral | 8 weeks | 8 weeks | ↑ tremors  ↔ choreoathetosis | (24) |
| Type I Pyrethroids (Pyrethrins, Resmethrin, S-bioallethrin, Permethrin, Bifenthrin, Tefluthrin) | Rat, Sprague Dawley, Male  n=10/group | Vehicle vs variable dosing per pyrethroid (low-dose with limited or no clinical signs vs high-dose with non-lethal, neurotoxic effects) for a single dose  Oral (gavage) | ~ PND 42 | ~ PND 42 | ↑ tremors, convulsions (moderate to severe)  ↑ sensory reflexes (exaggerated hindlimb flexion)  ↔ neuromuscular observations | (49) |
| Type II Pyrethroids  (Cypermethrin, Esfenvalerate, β-Cyfluthrin, Deltamethrin, Fenpropathrin, λ-Cyhalothrin) | Rat, Sprague Dawley, Male  n=10/group | Vehicle vs variable dosing per pyrethroid (low-dose with limited or no clinical signs vs high-dose with non-lethal, neurotoxic effects) for a single dose  Oral (gavage) | ~ PND 42 | ~ PND 42 | ↑ tremors, convulsions (slight)  ↑ sensory reflexes (strong)  ↑ neuromuscular observations (forelimb grip strength, reduced hindlimb grip strength, slower rotarod performance) | (49) |
| ***Neonicotinoids*** | | | | | | |
| Clothianidin | Mouse, C57BL/6N, Both sexes  n=6-10/group | Vehicle vs 80 mg/kg for a single dose  Oral (gavage) | 6 weeks | 3 or 7 months | ↑ tremors after administration  (did not last longer than 24 hours) | (10) |
| ***Organochlorines*** | | | | | | |
| Lindane | Rat, Wistar, Both sexes  n=4/sex/group | Vehicle vs 10 or 20 mg/kg for a single dose  Oral (gavage) | PD 22 | PD 22 | ↔ seizures  ↔ tremors | (44, 45) |
|  | Rat, Wistar, Both sexes  n=4/sex/group | Vehicle vs 10 or 20 mg/kg for a single dose  Oral (gavage) | PD 29 | PD 29 | ↔ seizures  ↔ tremors | (44, 45) |
|  | Rat, Wistar, Male  n=7-8/group | Vehicle vs 4 mg/kg for a single dose in animals prenatally exposed to saline or phencyclidine  **All animals in this study were exposed to lindane*  Intraperitoneal | PD 35 | PD 35 | ↑ convulsions | (50) |
